# Dual GLP-1 and GIP receptor agonist Tirzapetide plays an off-target role in the modulation of the β-adrenoceptors and glucose metabolism in hyperglycemic or senescent cardiac cells

**DOI:** 10.1101/2025.04.09.648025

**Authors:** Dunya Aydos, Zeynep Busra Aksoy, Mehmet Altay Unal, Kamil Can Akcali, Ceylan Verda Bitirim, Belma Turan

## Abstract

**Background:** A dual glucose-dependent insulinotropic polypeptide (GIP) and glucagon-like peptide-1 (GLP1) receptor agonist Tirzepatide (TZPD) is a novel cardioprotective agent, particularly in metabolic disturbances-related co-morbidities, however, there is no exact study to emphasize its possible off-target action in cardiac cells.

**Objective:** Taking into consideration a relationship between the trafficking of incretin receptors in a manner not anticipated by the standard way of cAMP as a primary actor in TZPD action, together with the role of cAMP depression in cardiac dysfunction, here, we aimed to elucidate a pattern of off-target receptor interactions of TZPD and molecular processes underlying the pleiotropic effects of TZPD through modulation of the β-adrenoceptors (β-ARs) signaling in cardiomyocytes.

**Methods:** To establish the multifaceted cardioprotective function and underlying mechanisms of TZPD against hyperglycemia (HG)- or senescence (SC)-induced cardiac dysfunction, H9c2 cells were treated with and without TZPD. We also used β_3_-ARs overexpressed H9c2 cells (β3OE) for comparisons.

**Results:** The TZPD intervention ameliorated the HG or SC phenotypes in the cardiac cells via alleviation in protein levels of GLP-1R and GIP-R as well as production of cAMP or cGMP even in the presence of these receptor antagonisms. The TZPD also alleviated the depressed levels of the β_1_- and β_2_-ARs with a significant decrease in the activated β_3_-ARs and PKG being parallel to normalizations in the cAMP and cGMP in the presence of the antagonisms of these receptors. The therapeutic effects of TZPD on the similar parameters of the β3OE group of cells can strongly verify its off-target action among multifaceted effects in either HG or SC cells. In addition, molecular dynamics simulations indicated that TZPD binds with the highest affinity to GLP-1R and β_3_-ARs rather than GIP-R and then relatively lower but almost similar affinities to β_1_- and β_2_-ARs. Furthermore, mechanistically, the cardioprotective effect of TZPD includes significant regulation of the cellular Ca^2+^, at most, modulating the proteins in β-ARs signaling pathways. Moreover, TZPD could significantly increase not only the depressed protein level but also the translocation of GLUT4 on the sarcolemma, promoting glucose uptake in the HG or SC groups independent of its receptor actions.

**Conclusions:** Our findings indicate that TZPD, with its multifaceted role, has beneficial effects on cardiac cells by positively modulating β-ARs signaling and glucose metabolism rather than on-target receptor action. Furthermore, we demonstrated how TZPD can engage the different targets with distinct signaling motifs at the sarcolemma.

**Highlights:**

- TZPD has direct cardio-therapeutic effects in cardiac cells under hyperglycemia or senescence, at most, through affecting altered β-ARs signaling in cardiomyocytes with the highest affinity to β_3_-ARs compared to the others.
- The multifaceted roles of TZPD in the HG or SC group of cells include modulation of β- ARs signaling, cellular Ca^2+^ regulation, and glucose metabolism independent from the insulin signaling pathway.
- TZPD could induce translocation of GLUT4 on the membrane and increase its protein level in the HG or SC group of cells independent of its receptor actions.
- TZDP could also normalize the depressed level of IRS-1in the HG or SC group of cells.
- TZPD activates both GLP-1R and GIP-R in cells, particularly with consideration of the in silico finding on the higher binding affinity of TZPD to GLP-1R rather than GIP-R, it seems an activation of GLP-1R by an agonist stimulates insulin secretion predominantly through the GLP-1R, with an additional contribution of GIP-R activation
- Overall, these results demonstrate that the same drug engaging the different targets has distinct signaling motifs at the plasma membrane and provides further information on the role of incretins in cardiac cells under hyperglycemia or senescence.

## Introduction

Diabetes is a chronic metabolic disease characterized by high blood glucose levels, which leads over time to serious damage to various organs including the heart, blood vessels, kidneys, and nerves. Epidemiological studies show the growing global burden for individuals, families, and countries for the prevalence of diabetes, most importantly, parallel to obesity. Obesity is a triggering factor for diabetes associated with insulin resistance, and therefore there are associations between type 2 diabetes, obesity, and insulin resistance (1). Interestingly, even early findings come up with an idea that there is a close relationship between age-related changes in insulin resistance and/or aging is a compulsory risk factor for insulin resistance which in turn plays an important role in cardiac aging (2, 3). Recent observations show that insulin can affect the activation of β-ARs and their signaling in the heart to modulate cardiac function in clinically relevant states (i.e. diabetes, heart failure, etc.), which in turn regulates cardiac glucose uptake (4). Insulin resistance also underlies a clustering of risk factors for cardiometabolic risk factors and even in the absence of systemic metabolic disorders, cardiac insulin resistance can develop various heart diseases (5, 6). Consequently, one can propose that there are crossroads between cardiac β-AR signaling and myocardial glucose metabolism in consideration of the interplay between diabetes, obesity, senescence, and insulin resistance (7). Therefore, a common treatment approach for metabolic disturbances associated with syndromes/diseases would be highly beneficial both economically and in implementation in clinics.

Correspondingly, anti-diabetic drugs, glucagon-like peptide-1 (GLP-1) agonists are recommended for treatment to reduce risks of major adverse cardiovascular events (8-10). Besides the known effect of GLP-1 in pancreatic cells, this incretin hormone provides action in the skeletal muscle independently from insulin receptor activation, while glucose-dependent insulinotropic polypeptide (GIP) exerts strong insulinotropic effects, however, its action on the muscle cells including cardiac cells remains to be examined (11, 12). Tirzepatide (TZPD) is a dual-receptor agonist that mimics two different hormones in biological systems, such as GLP-1 and GIP. In addition to its anti-diabetic effects, clinical studies emphasize its important cardioprotective effects not only in patients with T2DM but also in those with cardiac dysfunction via other pathological conditions (13-15). In this regard, authors have recently demonstrated that TZPD treatment significantly reduced the 10-year predicted risk of atherosclerotic cardiovascular disease compared to a placebo in nondiabetic patients with obesity or overweight, (16, 17). The GLP-1R and GIP-R are present in the heart of mammals and a dual incretin receptor agonist could present benefits by delivering superior effects through enhanced activation of insulin secretion and increases in intracellular cAMP in pancreatic β-cells (12, 18-22).

Moreover, at the pharmacodynamic level, TZPD has been identified as an imbalanced GLP-1R-GIPR co-agonist, implying a preference for GIP-R over GLP-1R with an action of GIP-R on reduction of GLP-1R internalization, and cAMP accumulation (19, 23-26). An interesting study by Chepurny and co-workers suggested that GIP-R or GLP-1R agonists may lead to off-target effects at other receptors by using molecular modeling analysis (27). In this aspect, others also pointed out a pattern of off-target receptor interactions, mostly at GPCRs, and demonstrated the structural probing of off-target GPCR activities within a series of agents (28, 29).

Previously published data mentioned that β_1_– and β_2_-ARs are either downregulated or desensitized, whereas the protein level of β_3_-ARs upregulates in mammalian heart under pathological condition, including insulin resistance (6, 30, 31). The β_1_– and β_2_-AR signaling pathways are both coupled to stimulation adenylyl cyclase (AC) and initiation of a transmembrane signal cascade which includes increases in intracellular cAMP, leading to the activation of cAMP-dependent PKA, while a much more efficiently coupled to the production of cAMP by stimulation of the β_2_-AR rather than the β_1_-AR (32, 33). On the other hand, the β_3_-AR expression is upregulated in the failing heart and there are important deleterious effects of β_3_□AR activation in cardiac remodeling under hyperglycemia or aging, in part, through a cross□link between β_3_□AR activation and NO□signaling pathways and inhibiting the production of cAMP (34-36). Recent condensed data pointed out the beneficial effects of not only GLP-1R agonists but also dual receptor agonists on the maintenance of heart function under pathological conditions (13-17, 27, 28), however, there are very limited documents to explain the underlying molecular mechanisms of regulation of cellular glucose metabolism in cardiac cells rather than insulin secretion. Taking into consideration the multifunctionality of GLP-1R as a vital component of the GPCR family and therapeutic target in various complex disease mechanisms (37), we hypothesize that dual receptor agonisms may have a direct off-target effect in cardiac cells to exert their beneficial effects on heart dysfunctions. Therefore, considering already known signaling pathways associated with cardiac dysfunction in the metabolic and/or elderly heart, along with the depression of cAMP levels (31), we investigated a possible off-target effect of TZPD in cardiomyocytes besides its dual receptor agonist action, through modulation of the β-ARs signaling and Ca^2+^ handling pathways. In addition, taking into consideration the stimulatory effect of GLP-1 on glucose transporters in β-cells (38), we also examined whether TZPD application includes an augmentation in GLUT4 trafficking in hyperglycemia- or senescence-mimicked cardiac cells.

## Methods and materials

### Cell line and treatment of cells

The H9c2 cell line was derived from the left ventricle of the embryonic rat heart (CRL1446). It was grown in modified Dulbecco’s modified Eagle’s medium (DMEM) (DMEM-LPA, Capricorn) including 5.5 mM glucose (low glucose) including 10% FBS (Gibco), 50 U/ml penicillin (Lonza, Swiss), 2 mM L-Glutamine (Lonza, Swiss), and 50 μg/mL streptomycin (Lonza, Swiss). To obtain hyperglycemic cells (HG-group), H9c2 cells were incubated with DMEM including 33 mM glucose for 48 h (Biowest, France). To obtain senescence (aging) model cells (SC-group), H9c2 cells were incubated with DMEM including 50 mg/mL D-galactose for 48 h. A β_3_-adrenergic receptor overexpression (β3OE group) in the H9c2 cell line using stable lentiviral infection (*see supplementary documents for details of the method*) and validation was performed as described previously (36). Untreated cells were kept as normal cells (NC group). The HG and SC groups of cultured H9c2 cells were treated with TZPD (GlpBio, GC3840) at concentrations ranging from 20 nM to 300 nM for 24 hours to assess cell viability To optimize the antagonist doses, the cultured HG and SC cells were incubated with GLP-1R Antagonist 1 (MedChem, HY-101116) at concentrations from 150 nM to 1.3 μM, and GIP-R Antagonist (MedChem, HY-P10138) at concentrations ranging from 25 nM to 240 nM for 24 hours. Cell viability was evaluated using the MTT [3-(4,5-dimethylthiazol-2-yl)-2,5-diphenyltetrazolium bromide] (Bioworld, 42000092) assay (*see supplementary documents for details of the method*). Note that antagonists and TZPD were administered in the same medium containing either high glucose or D-galactose. For the qRT-PCR and western blotting experiments, HG and SC cells were incubated with TZPD for 24 h; in other experiments, TZPD incubation was performed for 30 min.

### The mRNA level determination by qRT-PCR

To analyze mRNA levels, the total RNA of H9c2 cardiomyocytes was isolated by using RiboEx Reagent (GeneAll, 302-001), and the purified total-RNA was reverse transcribed with Entlink cDNA Synthesis-kit (Elk Biotech, E300), as described previously (39). A GoTaq® qPCR Master Mix (Promega, A6001) was used to quantify and amplify PCR products for each primer, and the primers’ specificity was controlled with known databases. The fold changes of genes were analyzed using the comparative (2^−ΔΔCt^) method. Cyclophilin was used as a housekeeping control for mRNA expression analysis as a housekeeping control. RT primers designed for each mRNA are given in Supplementary Table 1.

### Western blotting

The cells were gently washed two times with iced PBS and harvested at 4°C in lysis buffer containing 250 mM NaCl, 1% NP-40, and 50 mM Tris-HCl; pH 8.0 and 1XPIC. Total protein lysates were evaluated by BCA assay (Thermo Fisher, USA) and 30 µg protein from each group was loaded to 12,5% SDS-PAGE gel. Following the transfer and blocking steps with 1% BSA in TBS-0.3% Tween, the membrane was incubated with the primer antibodies of either β1-AR (Thermo,), β2-AR (Origine, NM_012492), β3-AR (Saint Jones, STJ91728-200), GLP-1R (MedChem, HY-P81160) GIP-R (MedChem, HY-P10138), PKG (Thermo, PA3-031A), Glut4 (Medchem, HY-P80689), pPKA (Cell signaling, 4781), PKA (Cell signaling, 4782), pGSK (Santa Cruz, sc-373800), GSK (Santa Cruz, sc-377213), eNOS (Thermo, PA3-031A), alpha tubulin (Santa Cruz, sc-5286), and GAPDH (Santa Cruz, sc-137179) primer antibody to use as house-keeping control. Anti-rabbit (Abcam, ab150077) and anti-goat (Santa Cruz, sc-2020) antibodies were used as seconder antibodies, respectively. Band intensities were calculated using Image J (NIH, USA). Band intensities were normalized according to GAPDH levels.

### Measurement of cAMP and cGMP levels

The cAMP level was measured by enzyme-linked immunoassay (ELISA) according to according to the manufacturer’s instructions (Cell Signaling, 4339S). Briefly, 50,000 cells/well were seeded in a 96-well plate. One group of the HG or SC cells was treated with antagonists for 30 min and TZPD for 30 min while the second group of them kept as untreated. Cells were then treated with 500 µM IBMX for 20 min and then flowed up a 10 µM forskolin application in the presence of IBMX for 10 min. Cells were then washed with 1xPBS and lysed for BCA assay. A protein sample with an amount of 200 µg was used from each group. A 50 μL sample was placed into the HRP-linked target solution and then transferred to the antibody-coated assay plate, letting it incubate for 3 h. Following the washing steps, the absorbance of the samples was determined at 450 nm.

In a similar procedure to cAMP determination, cGMP was also measured by Chemical’s Cyclic GMP ELISA Kit (Cayman, 581021). Control and treatment groups were lysed for BCA assay. Following the BCA assay, the enzymatic reaction procedure was performed according to the manufacturer’s instructions. Absorbance was measured at 450 nm.

### Determination of either GIP-R, GLP-1R, or GLUT4 localizations

Localizations of GIP-R, GLP-1R, and Glut4 in H9c2 cells were determined using anti-GIP-R (MedChem, HY-P10138), anti-GLP-1R (MedChem, HY-P81160) or anti-Glut4 (MedChem, HY-P80689) antibodies using confocal microscopy (Zeiss LSM 980). The plasma membrane was labeled using an E-cadherin monoclonal antibody (Biolegend, 866702). Here, we used the HG group of cells compared to the NC group. The loaded cells were treated with TZPD in the presence or absence of the GLP-1R and GIP-R antagonists for 30 min. To verify the localization of GLUT4, the NC group of cells was incubated with 100 nM insulin for 3 hours. After fixation and permeabilization of H9c2 cells with 4% paraformaldehyde and then 0.3% Triton-X100, the cells were incubated with primary antibody to monitor their localization on the membrane and cytoplasm. After overnight incubation of the cells, they were further incubated with appropriate secondary antibodies in the presence 0.25% Triton X-100/PBS for 5 min/each and incubated with Texas red goat anti-rabbit IgG (H&L) (1:500) (Rockland, 611-1902) and/or fluorescein goat anti-mouse IgG (H&L) (1:500) (Rockland, 610-1202) for 1 hour at room temperature. A mounting Medium with DAPI (Novus Biologicals, H-1200-NB) was used for mounting.

### Intracellular free Ca^2+^ determination in H9c2 cells

To detect the resting level of intracellular free Ca^2+^ in H9c2 cells, we loaded cells with a Ca^2+^-sensitive fluorescence indicator Fluo3-AM (MedChem, HY-126821). Then we used flow cytometry to detect the ratio of fluorescence changes (Acea, Novocyte). The trypsinized cells were incubated with 0.5 mM Fluo3-AM for 1 h in HBSS buffer. Here, we used the HG group of cells compared to the NC group. The loaded cells were treated with TZPD in the presence or absence of the GLP-1R and GIP-R antagonists for 30 min. Data were acquired with excitation of 488 nm wavelength.

### Glucose uptake assay

The glucose uptake content in the cells was determined in all groups of cells (i.e. HG group, SC group, NC group, β3OE) and calculated by the Glucose Uptake Fluorometric Assay Kit (Elabscience, E-BC-F041). The generated NADPH during the conversion of up-taken 2-DGbto 2-DG-6P by glucose dehydrogenase was detected and calculated by the fluorescence probe. The HG or the SC group of cells were treated with GIP-R and GLP-1R antagonists. Following 30 min of this group treatment, a 40 nM of TZPD was added into the medium and then incubated for 1 hour. After the incubation period, the cells were washed with Krebs– ringer phosphate HEPES (KRPH) buffer, and then 10 mmol/L 2-DG was added to the cell culture plates. Later, the cells were incubated at 37°C for 30 min. Following the treatment procedure, the supernatants were collected and placed in microplates, and a 50 μL sample was incubated with 250 μL enzyme at 37°C for 30 min.

Fluorescence intensity was measured at the excitation wavelength of 530 nm and the emission wavelength of 590 nm (Bertold). The proportion of glucose remaining in the supernatant was inversely correlated to the glucose taken to level up by the cells. The cells left in the cell culture as attached were also collected for BCA assay. BCA assay results were used for normalization.

### Statistics

Statistical analyses were performed using Prism 5.0 (GraphPad Software Inc.). For most experiments, three biologically independent replicates were performed and otherwise stated differently in the corresponding figure legend. The statistical difference between the two groups was analyzed using an unpaired Student’s T-test. Differences with two-tailed *p*-values□<□0.05 were considered statistically significant. All results are expressed as mean□±□standard error of the mean (SEM).

## Results

### Confirmation of expressions and localizations of GLP-1R and GIP-R in cardiac cells and their responses to TZPD application

Although the expression of GLP-1R has been shown in the H9c2 cell line, here we first investigated the localizations of either GIP-R or GLP-1R on the membrane of cardiac cells by using confocal laser scanning microscopy. The localization of these two receptors exists in normal control group (NC) cells with higher localizations on the membrane compared to the cytosolic part of the cells by using confocal microscopy (Fig. 1A and B, left parts). To examine the direct action of TZPD on the localizations of its receptors, we first performed the cell viability testing as responses of these cells to TZPD application (20-300 nM) by using the MTT test (Suppl. Fig. 1A). Second part of this group examinations, we prepared cells to mimic hyperglycemia (HG group; Fig. 1A and B; right parts) or senescence (SC group; Fig. 1A and B; middle parts) and then investigated the localizations of Texas red -tagged GLP-1R and Texas red -tagged GIP-R on the membrane vs. cytosol in the mimicked cells with and without TZPD application (40 nM). For note, the confirmation of the senescence mimic cell model (by D-galactose) was performed by using senescence-associated β-galactosidase (SA-β-Gal) testing in H9c2 cells (Supplementary Fig. 1B and C). As can be seen in Fig. 1A and B, the membrane localizations of these two receptors were markedly decreased in both the SC group (middle) and HG group (right) cells compared to those of the NC group of cells, TZDP treatment of these groups of cells provided marked recoveries in their membrane localizations.

**Fig. 1.**
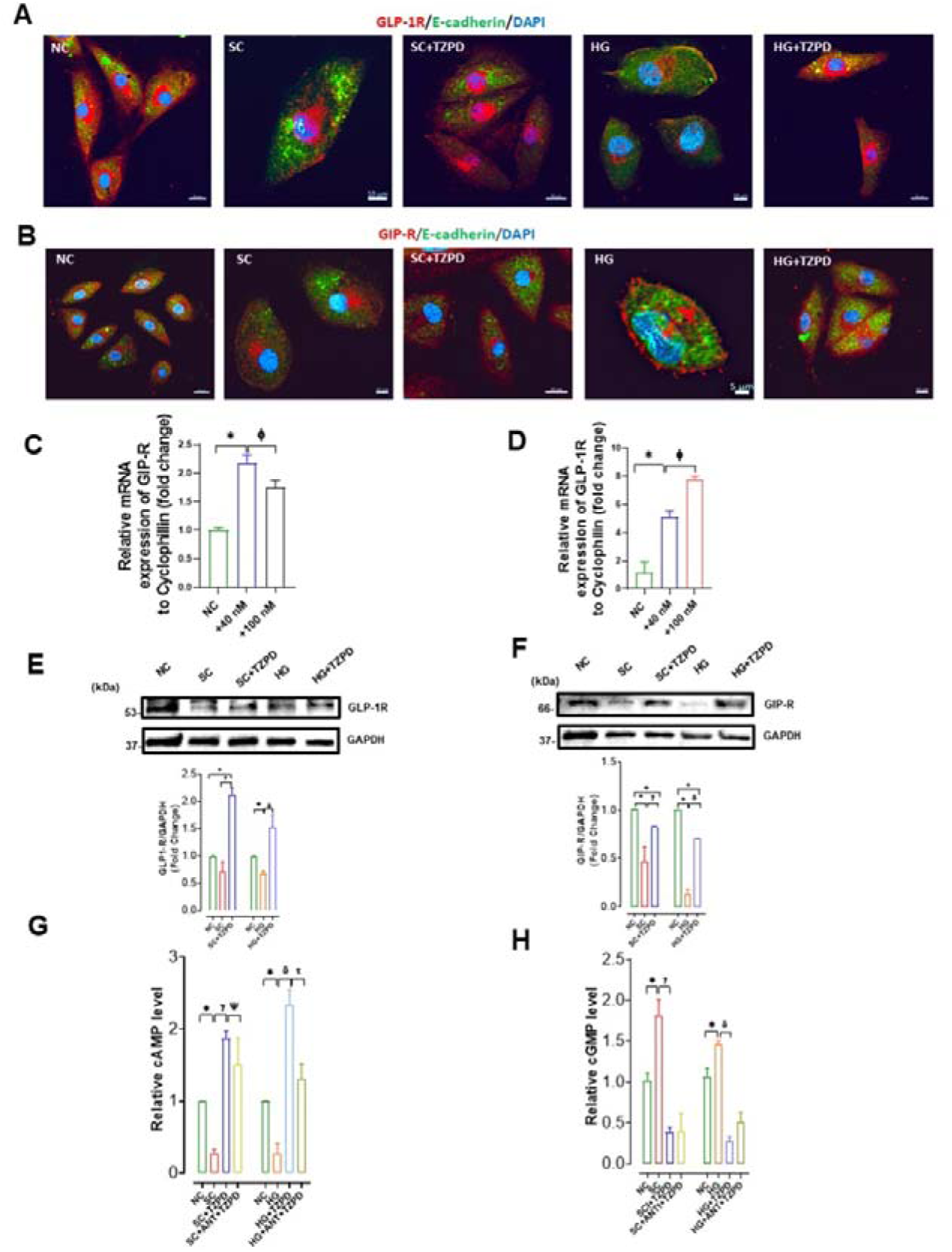
Demonstration of GLP-1R and GIP-R existence in H9c2 cells and their responses to Tirzepatide (TZPD) in a concentration-dependent manner. Verification of GLP-1R **(A)** of GIP-R **(B)** in immunostained H9c2 cells with either a Texas red -tagged GIP-R or a Texas red -tagged GLP-1R antibody. (NC group) by immunofluorescence microscopy. FITC-tagged E-cadherin was used to stain plasma membrane. The mRNA expression levels of GIP-R (**C**) or GLP-1R **(D)** with respect to Cyclophilin upon treatment with either 40 nM or 100 nM TZPD. TZPD application (40 nM for 24 h) provides significant augmentations in the depressed protein levels of GLP-1R **(E)** and GIP-R **(F)**, and the altered levels of cAMP **(G)** and cGMP **(H)** in either the senescence group of cells (SC group) or hyperglycemic group of cells (HG group) compared to the control group of cells (NC group). All shortens are the same in all figures after all. The protein levels were quantified by Western blotting and GAPDH levels were used for normalization. The representative protein bands are given in the upper parts of the bar graphs. The experimental groups are kept as two groups: SC+TZPD and SC+TZPD+ANT (under the presence of receptor antagonist) or HG+TZPD and HG+TZPD+ANT (under the presence of receptor antagonist). Figures include magnification=20X. Data are presented as means±SEM. \**p*<0.05 *vs.* NC group, ^φ^p<0.05 *vs.* +40 nM TZPD treated group, ^γ^*p*<0.05 *vs.* SC group, ^δ^*p*<0.05 *vs.* HG group, (*n*=3-4 independent experiments).

In the second part of these groups of examinations, we also determined the expression levels of GLP-1R and GIP-R by measuring mRNA levels under TZPD applications by using two different concentrations (24 h incubation). As can be seen in Fig 1(C and D, respectively), an application of 40 nM TZPD could induce about a 2.2-fold increase in the mRNA level of GIP-R while this increase was a 6-fold in the GLP-1R. When we increased the level of TZPD (100 nM), the response of GIP-R was decreased (about 25%), while the response of GLP-1R continued to increase (contrary to GIP-R) as about 8-fold compared to the TZPD untreated group of cells. These data showed the different responses to TZPD application in these two receptors. Therefore, following the above observation, the rest of all experimental investigations were performed by a 40 nM TZPD application by considering a similar level of response in these receptors with a TZPD application.

### Impact of TZPD on protein levels of GLP-1R and GIP-R as well as cAMP and cGMP production levels in HG- or SC-mimic cardiac cells

To examine the direct beneficial effect of TZPD on cardiac cells under HG or SC, we determined the protein levels of either GLP-1R (left) or GIP-R (right) by Western blotting in these mimicked cells compared to those of the normal cells (NC group). The protein levels of both GLP-1R and GIP-R were found to be significantly decreased under HG of SC conditions, and an addition of 40 nM of TZPD (24 h incubation) significantly prevented these decreases in their protein levels (Fig. 1E and F, respectively). Interestingly, the increase in the protein level of GLP-1R but not GIP-R following 40 nM TZPD application was even over its normal value determined in the NC cells. The effect of TZPD application on senescence- and aging-associated alterations was histologically confirmed through the β-Galactosidase and phosphorylated H2AX histone (termed γH2A.X) staining in HG- and SC-groups of cells (Supplement Fig. 1C). Our results demonstrated that TZPD treatment ameliorates SC and HG-associated senescence and DNA double-strand breaks.

Taking consideration the cAMP pathway in cardiomyocytes, particularly its relationship with β-adrenoceptors (β-ARs) signaling, to examine possible off-target effects of TZPD besides on-target effects in this signaling, with and without the existence of the receptor activities, we first examined the responses of GLP-1R and GIP-R to their antagonist applications by determination of cell viability (Supplement Fig. 2A, left and right, respectively). Taking into consideration a submaximal antagonist concentration for these receptors, we used antagonist concentrations as 650 nM for GLP-1R and 120 nM for GIP-R in our further examinations. Either a 650 nM antagonist (GLP-1R) application or a 120 nM antagonist (GIP-R) application could not significantly affect the level of cAMP in normal cells (NC group). Although there is no clear consensus on the status of either cAMP or cGMP levels in human failing *vs.* non-failing hearts (40), however, animal and/or cell-level studies mentioned important alterations in either plasma or cardiomyocyte levels of both cAMP and cGMP under hyperglycemia or senescence (6, 31). So, to verify whether TZPD can provide benefits on the parameters, we determined cAMP or cGMP levels in either the HG group or SC group of cells compared to the NC group of cells under 40 nM of TZPD application (24 h incubation). This application prevented significantly the decrease in cAMP levels in either the HG group or SC group, while increases in cGMP levels, even over their normal levels. Furthermore, these effects of TZPD are still almost similar levels under antagonist presences in these groups (slightly but significantly less compared to those without antagonists) (Fig. 1G and H, right and left, respectively).

### Verification of the off-target effect of TZPD in HG- or SC-associated alterations in cardiac cells through modulation of alterations in the β-ARs signaling

Previously, we and others have demonstrated significant changes in the protein levels of β- AR subgroups parallel to the changes in cAMP levels in the heart preparations under hyperglycemia, hyperinsulinemia, or aging (6, 31, 41, 42). Considering these relationships in cardiac cells between β-AR subgroups parallel to the changes in cAMP levels and/or cGMP levels, we first determined the responses to 40 nM TZPD application (24-h incubation) the protein levels of β_1_-, β_2_-, β_3_-AR in HG- or SC-groups of cells and then the responses to the TZPD application. The TZPD application (40 nM for 24 h incubation) could prevent the decreased protein level of β_1_-AR significantly with 100% prevention in the HG group of cells with about 75% recovery in the SC group of cells (Fig. 2A). The decreased protein level of β_2_-AR in the HG group was fully normalized in TZPD-applied cells (Fig. 2B), while the response to TZPD application induced a 2-fold increase in the protein level of SC group of cells compared to those of the NC group.

**Fig. 2.**
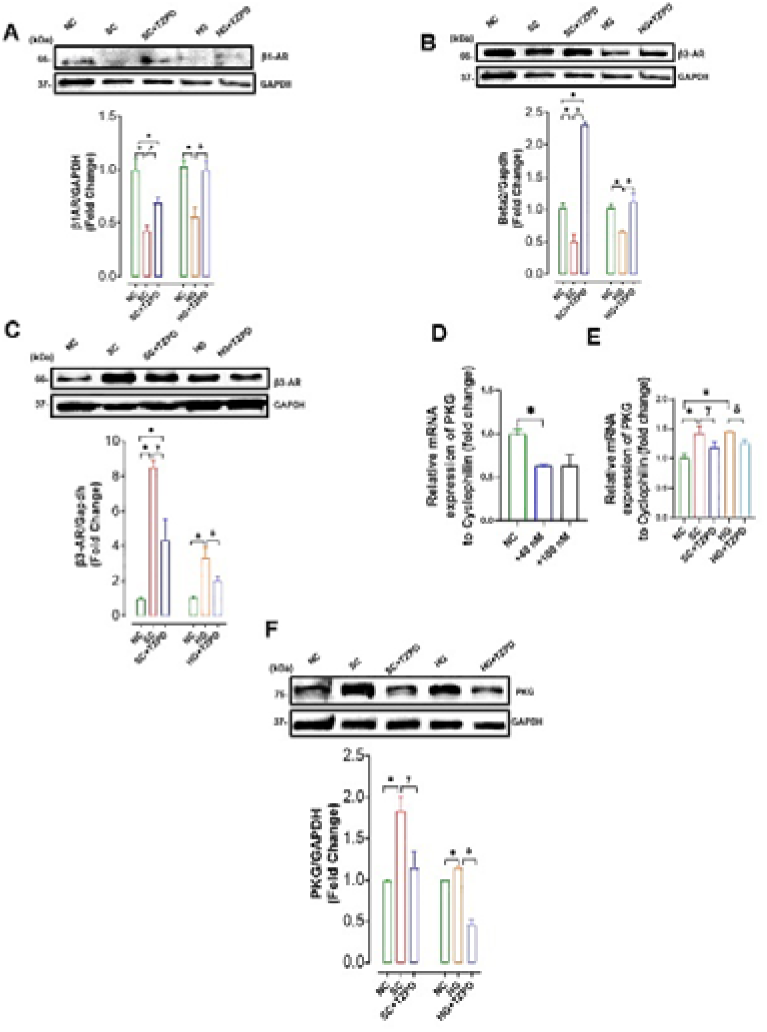
TZPD provides augmentations in the protein levels of β-AR systems as well as PKG of either the SC group or HG group H9c2 cells. The protein levels of β_1_-AR **(A)**, β_2_-AR **(B)**, and β_3_-AR **(C)** in either the SC group or HG group H9c2 cells treated with TZPD (40 nM for 24 h) compared to the NC group of cells. The TZPP with either 40 nM or 100 nM has significant inhibition of the mRNA level of PKG compared to the Cyclophillin the NC group of cells **(D)**. The TZPD application induced significant decreases in the activated mRNA levels of PKG in HG- or SC groups of cells compared to the NC group of cells **(E)**. **(F)** The 40 nM of TZPD application (24 h) provided a significant inhibition on the activated protein level of PKG in the SC group of cells (left) while it had much more inhibition on the protein level of PKG in the HG group of cells under its NC level (right). The protein levels were quantified by Western blotting and GAPDH levels were used for normalization. The representative protein bands are given in the upper parts of the bar graphs. Data are presented as means±SEM. \**p*<0.05 *vs.* NC group, ^φ^p<0.05 *vs.* +40 nM TZPD treated group, ^γ^*p*<0.05 *vs.* SC group, ^δ^*p*<0.05 *vs.* HG group.(*n*=3-4 independent experiments).

We also determined the protein levels of β_3_-AR in the HG group or SC group of cells under the TZPD application. As can be seen in Fig. 2C, the protein level of β_3_-AR was about 9-fold high in the SC group compared to the NC group while it was about 3.5-fold high in the HG group. The TZPD application to the cells (40 nM for 24 h incubation) could significantly prevent these increases in the protein level of β_3_-AR. When comparing the recoveries in the protein level of β-ARs by TZPD applications, there are significant differences depending on the subgroup of β-ARs as well as on the type of pathological condition (i.e. SC *vs* HG) in the cardiac cells.

### *In silico* analysis on the affinity of TZPD to either GLP-1R or GIP-R as well as β-ARs family

To confirm our experimental data on the effects of TZPD on either its receptors and/or other receptors/factors, we performed a series of in silico analyses to determine its affinity level to its receptors and possibly other receptors (Suppl. Fig. 3). To answer the first general question on what the interaction level of this agonist with the receptors, the energy level corresponding to the interaction level was calculated between Human GLP-1R protein and TZPD. Then it is tested to obtain the interaction whether or not with the same site as the 7FIM PDB ID structure, which is analyzed in the experimental structure analysis (the pink one). As seen in Suppl. Table 1, the interaction value in question is found to be -518.87 kcal/mol. The fact that this value is relatively low indicates that TZPD binds to this protein with high affinity. To a better level, exploring this topic is obtained by comparing the visual representations of the interactions. As can be seen in Suppl. Fig. 3, the interaction region (the pink one) (A) obtained after *in silico* analysis and the experimentally detected interaction region (B) are identical. Furthermore, the similar *in silico* analysis between Human GIP-R and TZPD presented an interaction value of -286.72 kcal/mol which is markedly lower than that of the previous interaction (Suppl. Fig. 3C).

We further repeated the similar *in silico* interaction analysis between TZPD and β_1_-AR, β_2_- AR, and β_3_-AR, respectively. As can be seen in Suppl. Table 2, the energy levels were found to be -274.81 kcal/mol, -263.55 kcal/mol, and -303.26 kcal/mol, respectively. In the framework of this proposition, since the interaction value of TZPD with β_3_-AR is closest to that of Human GLP-1R, the experimental results of our study design suggest that TZPD, at most, interacts with β_3_-AR protein with a very high affinity besides interaction with GLP-1R.

### The off-target effect of TZPD in HG- or SC-groups of cells includes the recovery in the level of PKG, a marker of activated β_3_-AR signaling

Since previous studies pointed out a relationship between a significant increase in the protein level of PKG, particularly in ventricular cardiomyocytes including substantial increases in the expression/protein level of β_3_-AR from either hyperglycemic heart or elderly heart of male rats (6, 36, 43), we first determined the responses of mRNA level of PKG to TZPD applications in NC group of cells. As presented in Fig. 2D, the mRNA level of PKG was found to be significantly decreased by either 40 nM or 100 nM of TZPD application. Then, we determined the responses of PKG to 40 nM of TZPD application in the HG or SC group of cells. As can be seen in Fig. 2E, the mRNA levels of PKG, in both the HG group and SC group of cells were significantly higher than those of the normal cells, while there were significant decreases in the mRNA levels of PKG following 40 nM of TZPD application.

We further determined the protein levels of PKG in either the HG group or SC group of cells and then the responses to 40 nM of TZPD application. The PKG protein level was found to be significantly high in either the HG group or the SC group, while there were significant recoveries in these cells following 24 h TZPD incubation (Fig. 2F). Interestingly, the decreased PKG protein level in the HG group was significantly less than that of the NC group. Of note, this observation implies a novel action (off-target action, independent from its receptors) of TZPD on activated β_3_-AR signaling in cardiomyocytes under pathological stimuli.

### Responses of the β-ARs family as well as cAMP and cGMP to TZPD in β_3_-AR overexpressed cardiac cells in the presence of GIP-R and GLP-1R antagonists in comparison in absence of antagonists

Taking into consideration the relationship between the activation of β_3_-AR, PKG, and the increase in cGMP in either the HG group or SC group, we determined the mRNA levels of either β_1_-AR, β_2_-AR, or β_3_-AR, and their responses to 40 nM of TZPD application in β_3_-AR overexpressed cells (β3OE). When compared to the without TZPD group, the mRNA levels of β_1_-AR and β_2_-AR in the β3OE group of cells were increased by about 1.5-fold and 4-fold by the TZPD application (40 nM, with 24 h incubation), respectively (Fig. 3A and B), whereas the mRNA level of β_3_-AR was found to decrease by about 10-fold (Fig. 3C).

**Fig. 3.**
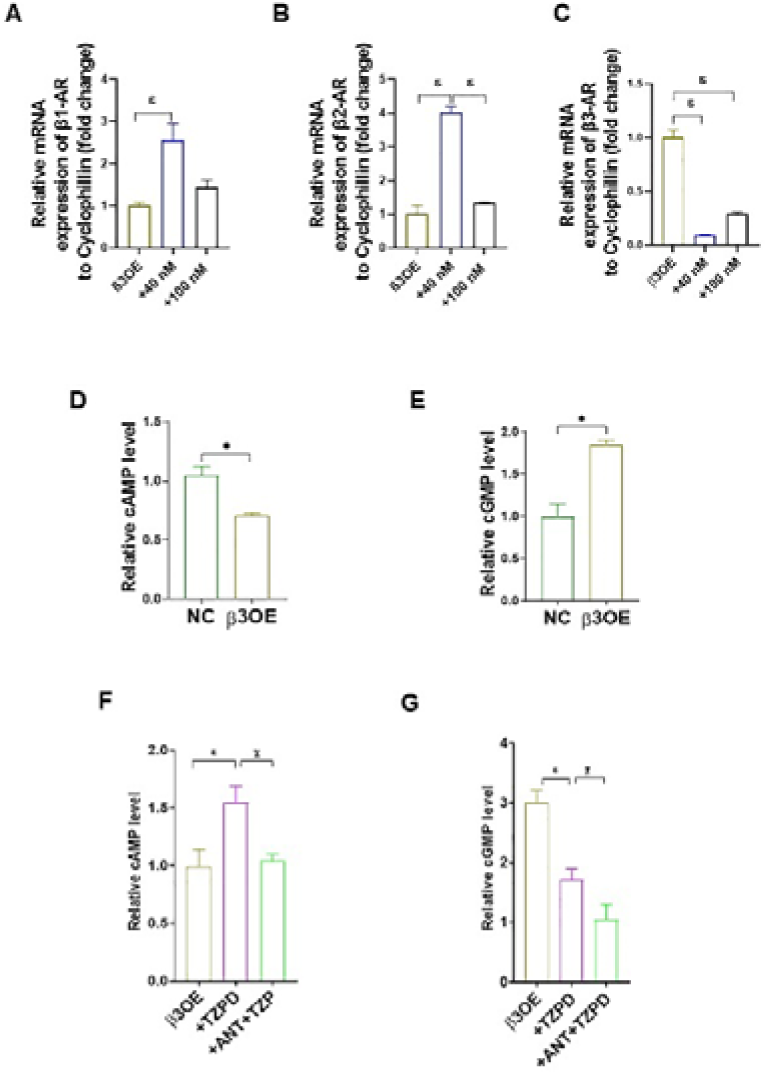
The TZPD application augmented the mRNA levels of β-ARs and the levels of cAMP and cGMP in β_3_-AR overexpressed (β3OE) H9c2 cells. The mRNA expression levels of β_1_-AR **(A)**, β_2_-AR **(B)**, and β_3_-AR **(C)** comparison to Cyclophillin in β3OE H9c2 cells upon treatment with 40 nM or 100 nM TZPD for 24 h. Cellular cAMP **(D)** and cGMP **(E)** levels in β3OE cells compared to those of the NC group of cells. Responses to the TZPD application in the production of cAMP **(F)** and cGMP **(G)** in β3OE cells with and without GLP-1R and GIP-R antagonisms. Data are presented as means±SEM. \**p*<0.05 *vs.* NC group, ^φ^p<0.05 *vs.* +40 nM TZPD treated group, ^ε^*p*<0.05 *vs.* β_3_OE group, ^χ^*p*<0.05 *vs.* +ANT+TZPD group (*n*=3-5 independent experiments).

When comparing the cAMP and cGMP levels in normal cells to those of the β3OE group of cells, the cAMP was about 25% less in the β3OE group, while the cGMP was about 75% higher in the same group of cells (Fig. 3D and E, respectively). In further examinations, we determined the responses of either cAMP or cGMP to the 40 nM of TZPD application in the β3OE group. The cAMP level of the β3OE group by the TZPD application, with the presence of antagonists of both receptors (GLP-1R and GIP-R) was found less compared to the without the presence of antagonists (Fig. 3F). Similarly, the cGMP level TZPD application under the same experimental protocols was found to be less compared to the without the presence of antagonists (Fig. 3G).

### Verification of the off-target effect of TZPD as the responses of mRNA of protein levels of GLP-1R, GIP-R, PKG, and eNOS in the β3OE group of cells

The responses to 40 nM of TZPD application (24 h incubation) of either GLP-1R or GIP-R as mRNA levels were also found to be significantly high in the β3OE group compared to the NC group (Fig. 4A and B, respectively). Interestingly, the response to the TZPD in the GLP-1R of this group of overexpressed cells was 10-fold higher, whereas this increase was 2.5-fold in the GLP-1R mRNA level. Further analysis showed that the mRNA level of PKG in the β3OE group of cells was found to be about 10-fold higher than those of the normal cells, while this high value was decreased to its half value following the TZPD application (Fig. 4C).

**Fig. 4.**
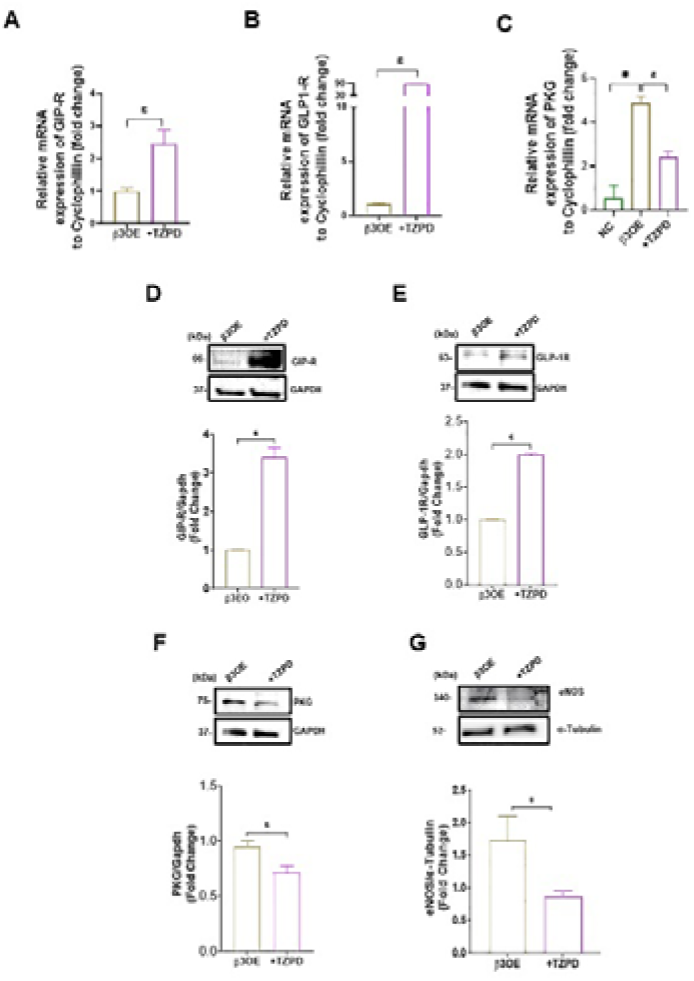
Effects of TZPD on mRNA and protein levels of GIP-R, GLP-1R, and PKG in β3OE cells. The responses to 40 nM or 100 nM of TZPD application as mRNA levels of GIP-R **(A)**, GLP-1R **(B)**, and PKG **(C)** compared to Cyclophill in β3OE cells. The protein levels of GIP-R, GLP-1R, and PKG under 40 nM of TZPD application in the β3OE cells compared to GAPDH analyzed by Western blot (D to F, respectively). The representative protein bands are given in the upper parts of the bar graphs. Data are presented as means±SEM. \**p*<0.05 *vs.* NC group, ^ε^*p*<0.05 *vs.* β_3_OE group (*n*=3 independent experiments).

Our Western blotting analysis of these receptors also confirmed that protein levels of either GLP-1R or GIP-R were significantly increased in the β3OE group of cells following 40 nM of TZPD application (Fig. 4D and E, respectively). Furthermore, when we examined the parameters associated with β_3_-AR overexpression which plays a critical role in regulating and maintaining a healthy cardiovascular system in our β3OE group of cells, the protein levels of PKG and total endothelial nitric oxide synthase (eNOS) were found to be decreased significantly in the same group of cells (Fig. 4F and G, respectively).

### TZPD recoveries cellular Ca^2+^ and marker proteins associated with Ca^2+^ handling in hyperglycemia-induced remodeling of cardiac cells

We first examined the basal level of Ca^2+^ in the HG-group of cells loaded with Fluo-3 dye compared to those of normal cells (NC-group) by flow cytometry. As shown in Fig. 5A, the level of Ca^2+^ is about 3.5-fold higher in the HG-group than in the NC group. Notably, the 40 nM of TZPD application with or without the receptor antagonists induced significant reversals.

**Fig. 5.**
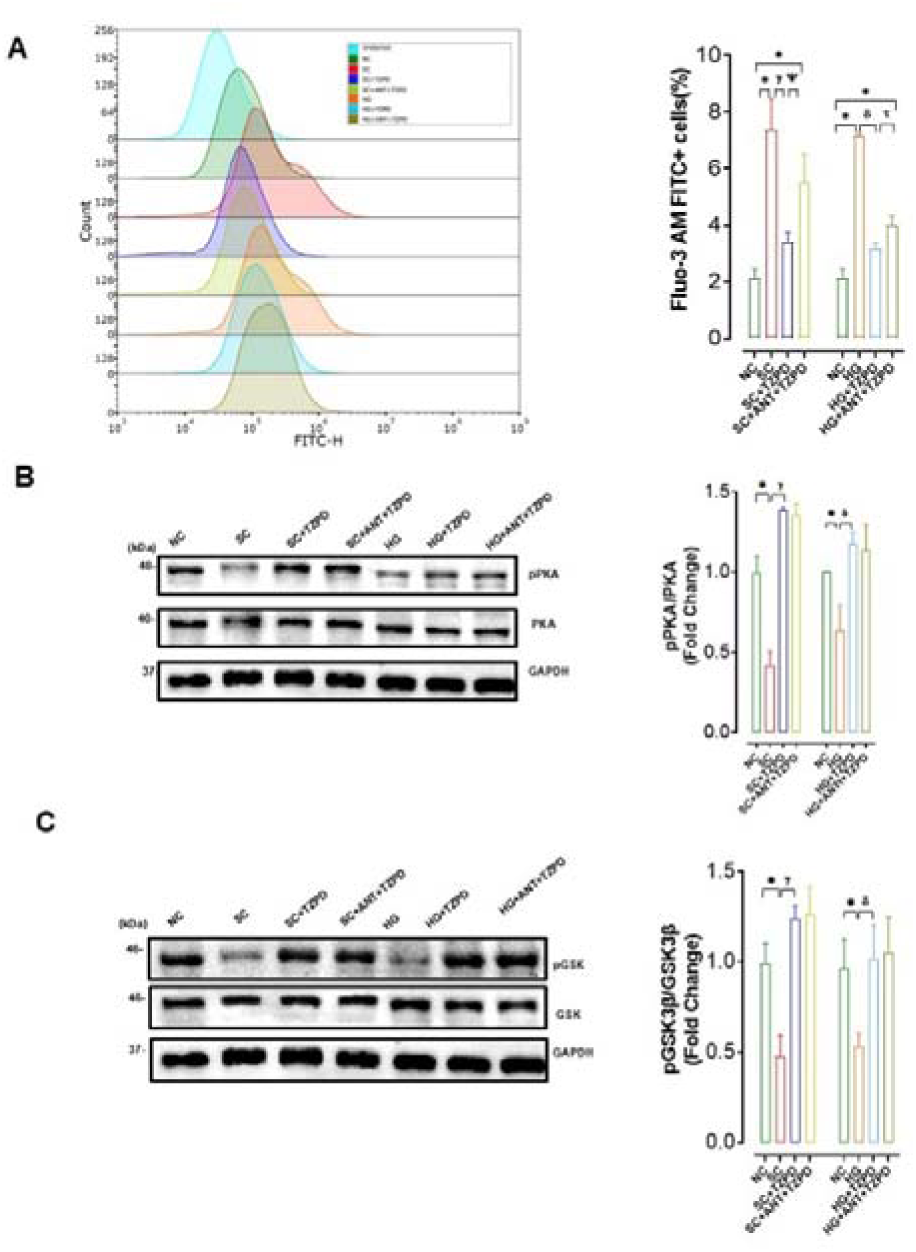
Effect of the TZPD application on the cellular resting level of Ca^2+^ and protein levels of parameters played roles in cellular Ca^2+^ regulation via β-AR signaling. **(A)** The cellular resting level of Ca^2+^ was imagined in all groups of H9c2 cells loaded with a Ca^2+^-sensitive fluorescence dye Fluo-3 AM. The representative images demonstrate the effect of TZPD (40 nM) on intracellular Ca^2+^ distribution in resting cells with and without receptor antagonism for the SC group of cells (left) and HG group of cells (right). The average cellular Ca^2+^ level in Fluo-3 AM loaded H9c2 cells was measured in all groups by the flow cytometer technique. Representative flow cytometry data are given in the left parts. The protein levels of phosphorylated PKA (pPKA) and PKA and their ratios **(B),** and phosphorylated GSK (pGSK) and GSK and their ratios **(C)**, compared to a reference protein GAPDH in the SC, SC+TZPD, SC+TZPD+ANT or HG, HG+TZPD, HG+TZPD+ANT comparison to the NC group. All shortens are the same as given in previous figures. Representative protein bands are given in the left parts of the bar graphs. Data are presented as means±SEM. \**p*<0.05 *vs.* NC group, ^γ^*p*<0.05 *vs.* SC group, ^δ^*p*<0.05 *vs.* HG group, ^Ψ^*p*<0.05 *vs.* SC+ANT+TZPD group, ^τ^*p*<0.05 *vs.* HG+ANT+TZPD group (*n*=3-4 independent experiments).

In addition, to support our electrophysiological data related to the cellular level of Ca^2+^, we examined the protein levels of some markers of cellular Ca^2+^ handling under the TZPD application to HG- or SC-cardiomyocytes. The 40 nM of the TZPD application significantly reserved the decreased protein levels of phosphorylated PKA (pPKA) without any significant effect on the PKA protein level in both SC (Fig. 5B). As can be seen in this figure, the decreased pPKA to PKA ratio is significantly increased to a level over its normal level. In addition, this increase was also similar to the increase under the presence of antagonists of GLP-1 and GIP receptors. On the left of the bar graphs, the representative protein bands are presented.

Another marker of cellular Ca^2+^ handling under TZPD application, we determined the protein level of phosphorylated GSK (pGSK) and GSK in either SC or HG groups of cells in comparison to those of NC cells. As can be seen in Fig. 5C, the pGSK/GSK was significantly depressed in both SC or HG groups of cells while TZPD application significantly recovered even over their normal values. The left parts in this figure show the representative protein levels. As an interesting observation similar to that of the previous one in (B), this increase was also similar to the increase under the presence of antagonists of GLP-1 and GIP receptors. On the left of the bar graphs, the representative protein bands are presented.

### TZPD increases protein levels and translocation of GLUT4 and promotes glucose uptake in the HG group cardiac cells or SC group cardiac cells

To investigate the role of TZPD on cellular glucose metabolism through GLUT4, we first imagined the GLUT4 distribution in cells and then quantified the distribution of GLUT4 as mRNA levels in the HG or SC groups of cells in comparison to the NC group (Fig. 6B). The representative original images are given in Fig. 6A. demonstrated the Texas-red tagged GLUT4 localization and FITC-tagged E-Cadherin staining.

**Fig. 6.**
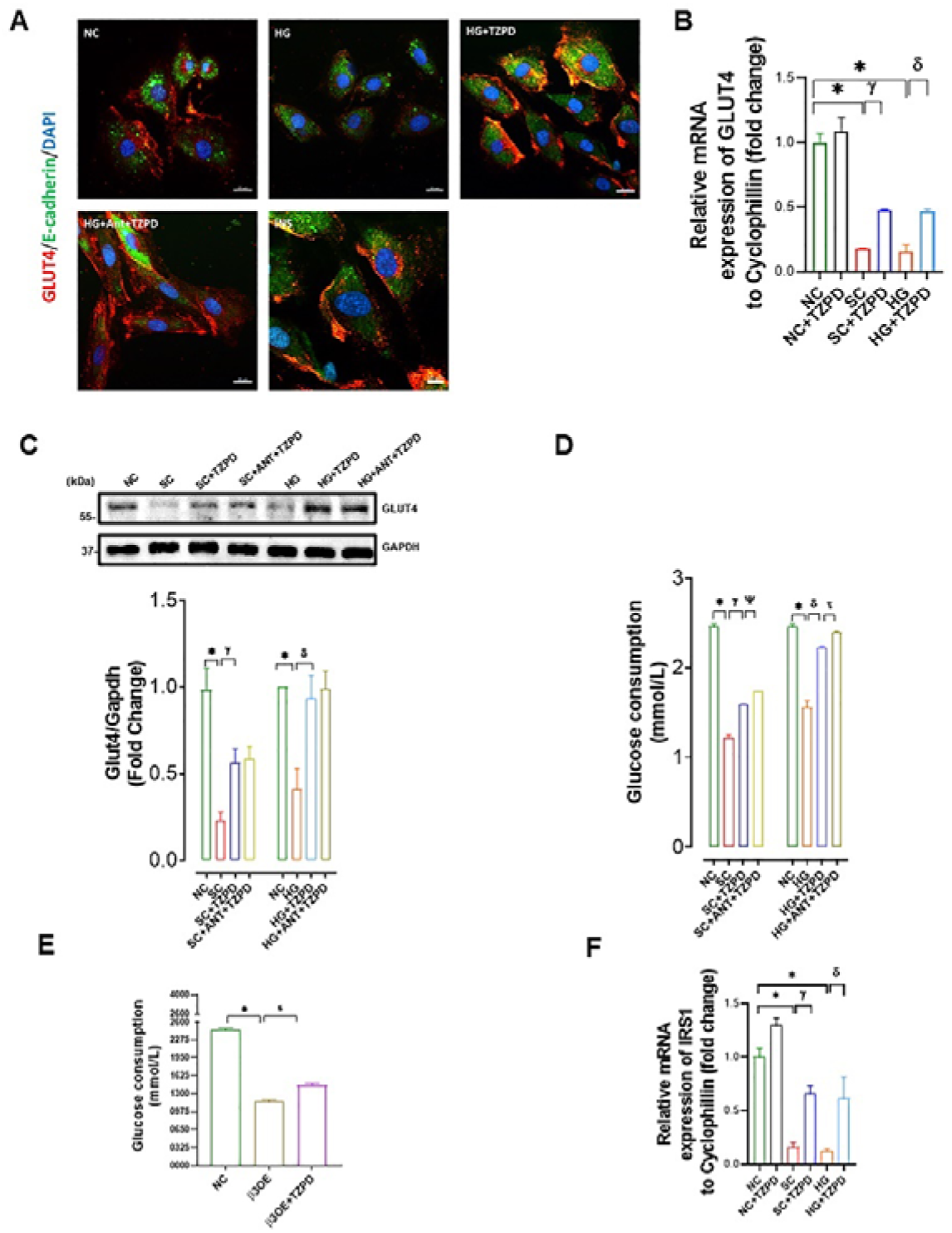
Effects of TZPD on cellular glucose uptake and translocation of GLUT4 on the cell membrane. **(A)** The distribution of GLUT4 in the cells when 40 nM of TZPD applied (30 min) to the SC group or HG group of cells in comparison to the NC cells given as representative confocal immunofluorescence staining and **(B)** the mRNA levels (relative to Cyclophillin) in these groups of cells when they incubated with 40 nM TZPD for 24 h. **(C)** The effects of the TZPD application on protein levels of GLUT4 in these experimental groups in comparison to the NC group with and without receptor antagonists. The protein levels were quantified by Western blotting and GAPDH levels were used for normalization. The representative protein bands are given in the upper parts of the bar graphs. **(D)** The levels of cellular glucose uptake in the SC (left) and the HG (right) groups of cells when 40 nM of TZPD applied (24 h) compared to those not applied with and without receptor antagonism and **(E)** the TZPD effect on glucose uptake in β3OE cells compared to those of NC group of cells. The mRNA level of IRS1 relative to Cyclophilin in experimental groups of cells with and without the TZPD application comparison to the NC cells **(F)**. GLUT4 imaging; 20X. Data are presented as means±SEM. \**p*<0.05 *vs.* NC group, ^γ^*p*<0.05 *vs.* SC group, ^δ^*p*<0.05 *vs.* HG group, ^Ψ^*p*<0.05 *vs.* SC+ANT+TZPD group, ^τ^*p*<0.05 *vs.* HG+ANT+TZPD group.(*n*=3-4 independent experiments).

Second, we determined the protein level of GLUT4 in the HG or SC groups of cells in comparison to the NC group under the 40 nM of TZPD application with or without the GLP-1R and GIP-R antagonists. As can be seen in Fig. 6C, GLUT4 protein levels in either the HG or the SC cells were found to be significantly decreased in comparison to the NC cells, while a 40 nM TZPD application (24 h incubation) provided a significant recovery in the protein levels. An interesting observation is to get a similar response to the TZPD under the presence of antagonists of GLP-1 and GIP receptors. The representative protein bands are presented on the upper parts of the bar graphs.

In the last part of this group investigation related to glucose metabolism, we established a glucose uptake process in the HG group of cells by TZPD application with and without receptor antagonist presence. We first used insulin to determine glucose uptake in the cells as the positive control (Suppl. Fig. 4). The cellular glucose consumption under insulin presence is maximum in the NC cells as expected. Also, the glucose uptake level of either the HG or the SC groups of cells was significantly less than that of the NC group cells, while 40 nM of TZPD application could recover significantly with and without the antagonists of both GLP-1R and GIP-R in a similar manner (Fig. 6D).

For further validation of the TZPD effect on cellular glucose mechanism, we determined the glucose uptake in the β3OE group of cells, as well. In comparison to that of the NC group, the uptake capacity of the β3OE group of cells was significantly lower, while it was recovered significantly by a 40 nM of TZPD application (Fig. 6E).

### Amelioration in IRS-1 level by TZPD can also confirm its off-target effects in the HG group or SC group

Since a decrease in insulin receptor substrate 1 (IRS-1) level is associated with remodeling in either hyperglycemic or elderly heart, we intended to examine whether TZPD affects another factor associated with cellular glucose metabolism in cardiac cells, particularly under hyperglycemic or senescence conditions, we first determined the mRNA levels of IRS1 following the TZPD application. As can be seen in Fig. 6F, TZPD application to the NS group of cells could induce a slight but significant increase in the mRNA level of IRS1. The mRNA level of IRN1 was a very high level of a significant decrease in both the HG and SC groups of cells, while the TZPD application significantly increased the depressed mRNA levels of IRS1 in both groups of cells. The recoveries in these levels were yet less than those of the NC group of cells level.

## Discussion

Our overall aim in the present study was to examine possible off-target effects of TZPD besides its dual receptor action in cardiac cells under hyperglycemia or senescence mimic conditions, thereby demonstrating that TZPD protects against diabetes-related heart damage of aging-related heart dysfunction in mammals. Here, our *in vitro* data supports the evidence of pleiotropic effects of TZPD which are mediated by modulation of β-AR signaling and cellular regulation of glucose metabolism together with Ca^2+^ homeostasis, at most, re-distribution of GLUT4 localization in the cardiac cells. So far, considering the clinal data on the beneficial effects of cardiac dysfunction under various pathological stimuli, our overall data provides important evidence of the TZPD effects in main molecular pathways involved in cardiac cells, particularly under hyperglycemia or senescence in mammal heart. Furthermore, our *in vitro* data strongly supports how/why TZPD application protects the heart against not only diabetes-related but also aging-associated damaging progresses in cardiomyocytes, at most, through modulation of β-ARs signaling and glucose metabolism, providing its direct multifaceted roles in cardiac cells.

The use of high-tech solutions in research and development offers numerous benefits, such as new drug development and their application fields, which unlock immense potential to optimize each stage of the process. Particularly, assuming new interactions for newly developed drugs is crucial for fully understanding their effects. Various approaches have been developed to infer potential interactions using bioactivity data, dose-response information, and large databases of known drug-target interactions (28, 44). Since a drug often can have off-target effects, however, under certain conditions, an off-target liability of a drug can contribute to therapeutic benefits by affecting different signaling pathways, primarily through binding, inhibition, degradation, or activation of receptors, rather than through its intended pharmacological target (45-47). In these regards, it is generally accepted that a dual incretin receptor agonist, TZPD has glucose-lowering in T2DM and anti-obesity effects through multiple molecular pathways, as well as protection against diabetes□related cardiac damages. The molecular mechanism of these beneficial effects of TZPD has been investigated in β-cells and interpreted through its biasing action at the GLP-1R and GIP-R to favor cAMP generation and then insulin secretion. In addition, some interesting pharmacological studies provided strong evidence that the same incretin group drugs (i.e. GLP-1 mimic GLP-1R agonists) can engage the different targets through distinct signaling motifs at the plasma membrane and provide further information related to the role of incretins in cardiac cells under pathological conditions (48). The studies on the comparison to drug binding kinetics, whether to its receptor or other membrane motifs, demonstrated their actions in a biased signaling of their trafficking profiles to improve agonist efficacy in various cell types except cardiomyocytes (37, 49, 50)

### TZPD application reverses the depressed protein levels of GLP-1 and GIP-R in the HG or SC group of cells without affecting their membrane localization

Incretin hormones including GIP and GLP-1 are important cellular metabolism regulators, at most, dependent on insulin signaling while their receptors are closely related members of the G-protein-coupled receptors (51-54). Various experimental data performed in animals and cell lines, mostly including pancreatic β-cells, demonstrated the existence of these receptors. Some of these studies demonstrated the mRNA levels of these receptors with less expression of GLP-1R than the GIP-R, particularly in cardiac cells including human AC16, mouse HL-1, and rat H9c2 (47, 55, 56). Here, we, first, confirmed the existence of GIP-R and GLP-1R together with their responses to TZPD in H9c2 cardiac cells and found significantly less expression of GLP-1 than the GIP-R in these cells like the previously published results in human AC16 cardiac cells (47). We have also confirmed the cellular and genetic response of H9c2 cells against TZPD in the HG and SC-mimicked groups. Increasing in β-Galactosidase and DNA double-strand breaks is considered one of the hallmark phenotypes of aged and hyperglycemic cells (57, 58). We demonstrated that TZPD treatment recovers the SC and HG-induced β-Gal and double-strand break formation.

Furthermore, we also examined the membrane localization of these two receptors by confocal microscopy in the first NC group and then the HG group while another group of investigation was performed in the SC group. Here, our reasoning for repeating the experiments with SC cells is the observation of the existence of insulin resistance in an important number of aged animals and their positive responses to insulin, similar to those of adult MetS animals (6, 43, 59). Besides, the freshly isolated cardiomyocytes isolated from both elderly rats and MetS rats have low cardiomyocyte cAMP levels, as well as higher cGMP levels in elderly cardiomyocytes than those of adult ones. These results are in line with our previously published data (6, 60). Therefore, here, being among our interesting findings, we, for the first time, have shown the role of TZPD in the SC group of cells through examinations of first the expressions and second localizations of GIP-R and GLP-1R.

As the first data in cardiac cells, our analysis demonstrated the membrane localizations of these two receptors are found to be similar in the HG or SC cells to those of the NC cells, while the TZPD application also could not affect the localizations of these two receptors in these cells. On the other hand, the protein levels of these two receptors are found to be depressed in both the HG and SC groups of cells compared to the NC cells. Correspondingly authors, taking into consideration the stimulation of insulin secretion by the incretin hormones diminished their levels in T2DM, particularly by hyperglycemia, examined the expression levels of GIP-R and GLP-1R and found to be significantly decreased levels while they were responding positively by insulin application (61). A further study in this content was performed in primary mouse islets and the β-cell line, and the findings demonstrated downregulation of GLP-1 receptor signaling by reducing its mRNA level and cell surface localization (54). Taken together these above results with our present data in cardiac cells, GLP-1R and GIP-R expression is decreased with chronic hyperglycemia, which further can confirm the relationship between depressed levels of these receptors and the impaired incretin effects found in T2DM. Under the light of already published data depending on either experimental findings and/or clinical outcomes, one can pronounce an interpretation of the relationship between reduced incretin hormone expressions together with loss of GLP-1R from the cell surface under hyperglycemia, which may correspond to the reduced clinical efficacy of GLP-1R based therapies in T2DM patients. This interpretation is further supported by the study of Willard et al. who demonstrated that TZPD induces differential internalization of the GIP-R and GLP-1R (23). Experiments in primary islets revealed a conclusion on the biased agonism of TZPD including enhancement of insulin secretion. Consequently, it seems there are important combinations of distinct signaling properties to the biased agonism of TZPD, which further may account for the promising efficacy of TZPD in damaged cells including cardiomyocytes.

The responses of GLP-1R to TZPD in the HG and SC groups of cells were found to be much higher even over control values than those of the GIP-R under similar experimental conditions. Although TZPD is a dual receptor incretin agonist and mediates insulin responses in β-cells, the GLP-1R is a drug target for the treatment of T2DM whereas the therapeutic potential of alone GIP-R is not very well defined. However, since TZPD activates both GLP-1R and GIP-R in cell lines and animal models, particularly with consideration of the in silico finding on the higher binding affinity of TZPD to GLP-1R rather than GIP-R, it seems an activation of GLP-1R by an agonist stimulates insulin secretion predominantly through the GLP-1R, with an additional contribution of GIP-R activation (62, 63).

### A positive effect of TZPD on the production of cAMP and cGMP levels in the HG or SC group of cardiac cells seems to depend on not only its receptor mediation but also a combination effect associated with mediation of β-ARs signaling

Various clinical data but less experimental findings announced that a dual incretin receptor agonist, TZPD, has potential cardiovascular benefits under many types of pathological conditions besides its glucose-lowering and anti-obesity effects. However, some experimental findings pointed out its integrated potency and signaling properties to provide a broad metabolic control mimicking the actions of native GIP and GLP-1 in the mammalian body. As already mentioned in previous findings, the positive effect of TZPD has been demonstrated through its biasing action at the GLP-1R comparison to GIP-R to favor AC-mediated cAMP elevation and subsequent activation of cAMP-dependent protein kinase(s) which stimulates the insulin release in β-cells. In addition, it is reported that GIPR agonism improves insulin sensitivity by a mechanism independent of glucose level or weight loss (62). Moreover, experimental evidence has revealed that GLP-1 and its analogs possess cardioprotective effects by various mechanisms related to cardiac contractility, myocardial glucose uptake, cardiac oxidative stress and ischemia/reperfusion injury, and mitochondrial homeostasis (59). Considering the general knowledge about compensatory and adaptive changes that occur in the heart to preserve cardiac output, supporting studies demonstrated that GLP-1R signaling activation by agonism can activate also different close targets inducing either adverse drug reactions or beneficial actions through different receptor-mediated signaling pathways (37). Using recent biosensors, authors explored the ability of GLP-1R activation by agonists in turn activating various distinct plasma membrane signaling profiles, directly or indirectly, thereby, providing improvements to the therapeutic potential of GLP-1R agonists.

In this context, the β-ARs belong to the superfamily of membrane proteins known as G-protein-coupled receptors (GPCRs). Under physiological conditions, cAMP signaling plays a key role in the regulation of cardiac function. The activation of this intracellular signaling pathway mirrors cardiomyocyte adaptation to various extracellular stimuli. Extracellular ligand binding to seven transmembrane receptors (also known as GPCRs) with G proteins and ACs modulate the intracellular cAMP content. Subsequently, this second messenger triggers the activation of specific intracellular downstream effectors that ensure a proper cellular response. Therefore, the cell needs to keep the cAMP signaling highly regulated in space and time.

### *In silico* analysis confirms the off-target action of TZPD through modification in the β-ARs family

A dual GIP/GLP-1 receptor agonist TZPD having the form of a synthetic linear peptide, contains 39 amino acids conjugated with the C20 fatty acid molecule which has a linker to a lysine residue at a position of 20 and the sequence of TZPD peptide also contains two non-coding amino acid residues. Studies demonstrated that TZPD binds to GIP-R and GLP-1R with a 5-fold weaker affinity than native GLP-1 for GLP-1R (64). In addition, pharmacological studies on cellular signaling processes demonstrated the TZPD as an imbalanced agonist of the GIP-R and GLP-1R (23). By using pancreatic β-cells, it has been shown that these two receptors can show biased signaling at the GLP-1R to favor cAMP generation, providing evidence of a weaker ability to drive GLP-1 receptor internalization compared with natural GLP-1. Furthermore, investigations on whether GLP-1R agonists with distinct signaling profiles demonstrated that there are possible 15 different pathways at the plasma membrane by systematically examining their plasma membrane to cytosolic localizations (37). Consequently, these types of analysis can provide important information on the interactions of GLP-1R agonists with other membrane receptors/kinases, including G protein-coupled receptor kinases (GRKs). Considering the general fact associated with cardiac function and mainly controlled by β-ARs signaling, which is also controlled by GRKs, our hypothesis on the off-target action of TZPD in cardiac cells may include β-Ars and their signaling pathways seems perfectly to be confirmed by our current data including in silico analysis. The comparison of the calculated energy levels for TZPD interaction of either its receptors or β-AR family members demonstrated that TZPD interaction has a higher affinity to GLP-1R than GIP-R, which increases their protein levels but not membrane localizations. However, this drug has its beneficial effect by affecting the internalization of these receptors and consequently release of insulin in pancreatic β-cells (19, 25). Considering the cardiac cells do not have any insulin vesicles and no insulin release, one can propose that TZPD has different beneficial effects in cardiac cells, at most, through inducing beneficial modulations in β-AR family members and their signaling pathways, being independent of a direct insulin signaling.

### TZPD application contributes directly to glucose metabolism in hyperglycemic cardiac cells through the translocation of GLUT4 into the plasma membrane

Recent studies, both *in vitro* and *in vivo* showed that cardiac β-AR signaling has an important impact on myocardial glucose metabolism, independent of insulin signaling but through regulation of GLUT4 into the membrane and thereby glucose uptake by myocytes (65). In this pathway, findings strongly supported the role of activation and protein levels of β-ARs, particularly, β_2_-AR associated increases in translocation of GLUT4 via GPCR kinase sites (33). According to our general knowledge, heart muscles have high glucose consumption, and glucose transport into cardiomyocytes falls behind the need if there is either an absence of insulin or less plasma membrane GLUT4 localization.

In this context, our findings showed that TZPD application in cardiac cells significantly increased the glucose uptake in both HG and SC groups of cells. Indeed, our present data being parallel to previously published demonstrated a marked decrease in glucose uptake (). By considering the role of GLUT4 membrane localization in the glucose uptake, our observation on this re-localization of GLUT4 under TZPD application can be explained by the recoveries in β-ARs, including of β_1_-ARs and β_2_-ARs, at most via PKA. In addition, the significant interaction between TZPD and these β-ARs by *in silico* analysis can imply the possible role of these activation/interactions in β-ARs to GLUT4 localization into the plasma membrane.

Interestingly, our findings also demonstrated that the inhibited expression of IRS-1 in either the HG or SC groups of cells could be recovered by TZPD application. This recovery also can be another source for the translocation of GLUT4 into the plasma membrane. Consequently, dual receptor agonist TZPD, being independent of insulin secretive action in β-cells, here, can present an important action on the regulation of cardiac cell glucose metabolism.

### The beneficial effect of TZPD in the pathologic heart including particularly its off-target action on β_3_-ARs can strongly support the statement of β_3_-ARs as new therapeutic targets for cardiovascular pathologies

The electrical activity of the heart is regulated by a sympathetic nervous system which is also mediated by β-ARs. The three major cardiac β-AR subtypes, β_1_- and β_2_-ARs, mediate intracellular signaling pathways regulating contractility via a positive inotropic effect of catecholamines through activation of AC-associated cAMP production under physiological conditions, whereas the activation of β_3_-ARs in the remodeling of the heart under metabolic disturbances (i.e. diabetic cardiomyopathy, metabolic heart, elderly heart, etc.) exerts a negatively inotropic effect through activation of NO signaling pathway including activation of eNOS and PKG (31, 36, 42, 66-70). Our cells mimicking either hyperglycemia or elderly mammalian heart cardiomyocytes have responded to TZPD application in terms of β-AR responsiveness. Not only the depressed protein levels of both β_1_- and β_2_-ARs but also their contribution to cAMP production significantly accelerated. In addition, the highly expressed β_3_-ARs and its associated signaling factors such as PKG and cGMP in these groups of cells were significantly recovered by TZPD application. More importantly, the similar beneficial effects of TZPD were also determined in the β3OE group of cells. More importantly, the energy level calculation for the interaction of TZPD with β_3_-ARs strongly supports our interpretation. Overall, one can sum up that TZPD, has an important off-target action in the β-AR system in cardiac cells under metabolic disturbances in the heart, at most, through mediating the regulation of β_3_-ARs signaling and therefore, provides an important indication about the TZPD application as a new therapeutic agent against new therapeutic targets for cardiovascular pathologies.

## Conclusions

In the present study, consistent with our previous results, TZPD has important beneficial effects on the metabolic heart, through regulation of cardiac function and glucose metabolism. The TZPD-associated recovery in high resting levels of free Ca^2+^ in the HG or SC groups of cells together with recoveries in the phosphorylation/protein levels of signaling proteins play critical roles in the Ca^2+^ regulation. Overall, our data, for the first time performed in cardiac cells, demonstrated that TZPD has an off-target effect in metabolic heart or elderly heart through modulation of altered β-AR system signaling and cellular free Ca^2+^ regulation besides receptor-related effect. In addition, it can regulate the glucose metabolism in cardiac cells, in a manner independent of its receptor action (Fig. 7).

**Fig. 7.**
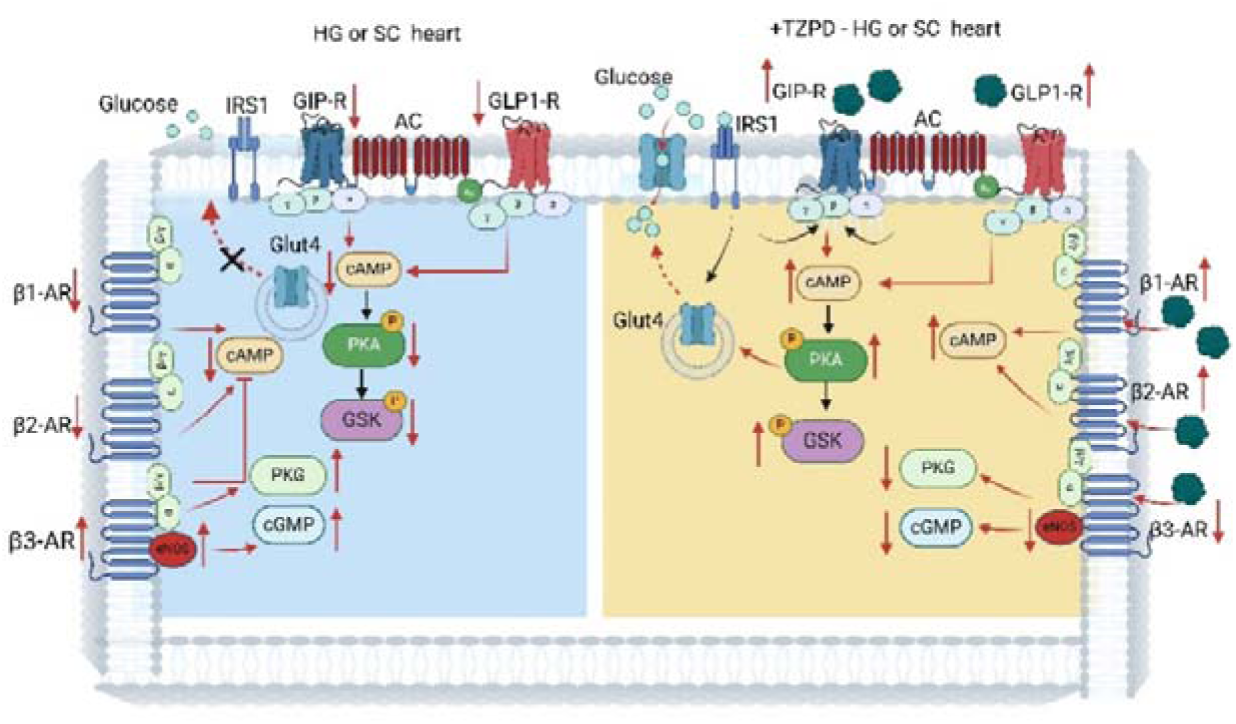
Schematic representation of our hypothesis and the findings in cardiac cells by application of TZPD. Our present data are accepted as an off-target effect in metabolic or elderly heart through modulation of altered β-AR system signaling and cellular free Ca^2+^ regulation besides receptor-related effect. In addition, the TZPD directly affected the regulation of the glucose metabolism in cardiac cells, in a manner independent of its receptor action.

## Acknowledgment

The authors would like to thank Dr. Y. Olgar for his technical support.

## Author Contributions

All authors have approved this manuscript and its contents, and they are aware of the responsibilities connected with authorship. Belma Turan and Ceylan Verda Bitirim contributed to the conception and design of the study and reviewed and approved the final version of all data. Kamil Can Akcali reviewed and approved the final version of the manuscript. Dunya Aydos, and Zeynep Busra Aksoy performed the majority of the experiments and their analysis. Mehmet Altay Unal contributed to the *in silico* analysis of the drug.

## REFERENCES

1. Wondmkun YT. Obesity, Insulin Resistance, and Type 2 Diabetes: Associations and Therapeutic Implications. Diabetes Metab Syndr Obes. 2020;13:3611–6.

2. Ilkun O, Boudina S. Cardiac dysfunction and oxidative stress in the metabolic syndrome: an update on antioxidant therapies. Curr Pharm Des. 2013;19(27):4806–17.

3. Rizvi AA, Rizzo M. Age-Related Changes in Insulin Resistance and Muscle Mass: Clinical Implications in Obese Older Adults. Medicina (Kaunas). 2024;60(10).

4. Fu Q, Wang Q, Xiang YK. Insulin and beta Adrenergic Receptor Signaling: Crosstalk in Heart. Trends in endocrinology and metabolism: TEM. 2017;28(6):416–27.

5. Adeva-Andany MM, Martinez-Rodriguez J, Gonzalez-Lucan M, Fernandez-Fernandez C, Castro-Quintela E. Insulin resistance is a cardiovascular risk factor in humans. Diabetes Metab Syndr. 2019;13(2):1449–55.

6. Olgar Y, Durak A, Bitirim CV, Tuncay E, Turan B. Insulin acts as an atypical KCNQ1/KCNE1-current activator and reverses long QT in insulin-resistant aged rats by accelerating the ventricular action potential repolarization through affecting the beta(3) - adrenergic receptor signaling pathway. J Cell Physiol. 2022;237(2):1353–71.

7. Shimi G, Sohouli MH, Ghorbani A, Shakery A, Zand H. The interplay between obesity, immunosenescence, and insulin resistance. Immun Ageing. 2024;21(1):13.

8. Marso SP, Bain SC, Consoli A, Eliaschewitz FG, Jodar E, Leiter LA, et al. Semaglutide and Cardiovascular Outcomes in Patients with Type 2 Diabetes. N Engl J Med. 2016;375(19):1834–44.

9. Gerstein HC, Colhoun HM, Dagenais GR, Diaz R, Lakshmanan M, Pais P, et al. Dulaglutide and cardiovascular outcomes in type 2 diabetes (REWIND): a double-blind, randomised placebo-controlled trial. Lancet. 2019;394(10193):121–30.

10. Piccini S, Favacchio G, Panico C, Morenghi E, Folli F, Mazziotti G, et al. Time-dependent effect of GLP-1 receptor agonists on cardiovascular benefits: a real-world study. Cardiovasc Diabetol. 2023;22(1):69.

11. Holst JJ. The physiology of glucagon-like peptide 1. Physiol Rev. 2007;87(4):1409–39.

12. Seino Y, Fukushima M, Yabe D. GIP and GLP-1, the two incretin hormones: Similarities and differences. J Diabetes Investig. 2010;1(1-2):8–23.

13. Nicholls SJ, Nelson AJ, Ditmarsch M, Kastelein JJP, Ballantyne CM, Ray KK, et al. Obicetrapib on top of maximally tolerated lipid-modifying therapies in participants with or at high risk for atherosclerotic cardiovascular disease: rationale and designs of BROADWAY and BROOKLYN. Am Heart J. 2024;274:32–45.

14. Lingvay I, Mosenzon O, Brown K, Cui X, O’Neill C, Fernandez Lando L, et al. Systolic blood pressure reduction with tirzepatide in patients with type 2 diabetes: insights from SURPASS clinical program. Cardiovasc Diabetol. 2023;22(1):66.

15. Nauck MA, D’Alessio DA. Tirzepatide, a dual GIP/GLP-1 receptor co-agonist for the treatment of type 2 diabetes with unmatched effectiveness regrading glycaemic control and body weight reduction. Cardiovasc Diabetol. 2022;21(1):169.

16. Hankosky ER, Wang H, Neff LM, Kan H, Wang F, Ahmad NN, et al. Tirzepatide reduces the predicted risk of developing type 2 diabetes in people with obesity or overweight: Post hoc analysis of the SURMOUNT-1 trial. Diabetes Obes Metab. 2023;25(12):3748–56.

17. Mori Y, Matsui T, Hirano T, Yamagishi SI. GIP as a Potential Therapeutic Target for Atherosclerotic Cardiovascular Disease-A Systematic Review. Int J Mol Sci. 2020;21(4).

18. Aguilar-Ballester M, Herrero-Cervera A, Vinue A, Martinez-Hervas S, Gonzalez-Navarro H. Impact of Cholesterol Metabolism in Immune Cell Function and Atherosclerosis. Nutrients. 2020;12(7).

19. Nuamnaichati N, Mangmool S, Chattipakorn N, Parichatikanond W. Stimulation of GLP-1 Receptor Inhibits Methylglyoxal-Induced Mitochondrial Dysfunctions in H9c2 Cardiomyoblasts: Potential Role of Epac/PI3K/Akt Pathway. Front Pharmacol. 2020;11:805.

20. Tian X, Gao Y, Kong M, Zhao L, Xing E, Sun Q, et al. GLP□1 receptor agonist protects palmitate-induced insulin resistance in skeletal muscle cells by up-regulating sestrin2 to promote autophagy. Sci Rep. 2023;13(1):9446.

21. Mayendraraj A, Rosenkilde MM, Gasbjerg LS. GLP-1 and GIP receptor signaling in beta cells - A review of receptor interactions and co-stimulation. Peptides. 2022;151:170749.

22. Kang MG, Campbell KP. Gamma subunit of voltage-activated calcium channels. The Journal of biological chemistry. 2003;278(24):21315–8.

23. Willard FS, Douros JD, Gabe MB, Showalter AD, Wainscott DB, Suter TM, et al. Tirzepatide is an imbalanced and biased dual GIP and GLP-1 receptor agonist. JCI Insight. 2020;5(17).

24. Hiromura M, Mori Y, Kohashi K, Terasaki M, Shinmura K, Negoro T, et al. Suppressive Effects of Glucose-Dependent Insulinotropic Polypeptide on Cardiac Hypertrophy and Fibrosis in Angiotensin II-Infused Mouse Models. Circ J. 2016;80(9):1988–97.

25. Kuna RS, Girada SB, Asalla S, Vallentyne J, Maddika S, Patterson JT, et al. Glucagon-like peptide-1 receptor-mediated endosomal cAMP generation promotes glucose-stimulated insulin secretion in pancreatic beta-cells. Am J Physiol Endocrinol Metab. 2013;305(2):E161–70.

26. Roed SN, Nohr AC, Wismann P, Iversen H, Brauner-Osborne H, Knudsen SM, et al. Functional consequences of glucagon-like peptide-1 receptor cross-talk and trafficking. The Journal of biological chemistry. 2015;290(2):1233–43.

27. Chepurny OG, Matsoukas MT, Liapakis G, Leech CA, Milliken BT, Doyle RP, et al. Nonconventional glucagon and GLP-1 receptor agonist and antagonist interplay at the GLP-1 receptor revealed in high-throughput FRET assays for cAMP. The Journal of biological chemistry. 2019;294(10):3514–31.

28. Meimetis N, Lauffenburger DA, Nilsson A. Protocol to infer off-target effects of drugs on cellular signaling using interactome-based deep learning. STAR Protoc. 2025;6(1):103573.

29. Paoletta S, Tosh DK, Salvemini D, Jacobson KA. Structural probing of off-target G protein-coupled receptor activities within a series of adenosine/adenine congeners. PloS one. 2014;9(5):e97858.

30. Brodde OE, Michel MC. Adrenergic and muscarinic receptors in the human heart. Pharmacol Rev. 1999;51(4):651–90.

31. Okatan EN, Tuncay E, Hafez G, Turan B. Profiling of cardiac beta-adrenoceptor subtypes in the cardiac left ventricle of rats with metabolic syndrome: Comparison with streptozotocin-induced diabetic rats. Can J Physiol Pharmacol. 2015;93(7):517–25.

32. Liang W, Curran PK, Hoang Q, Moreland RT, Fishman PH. Differences in endosomal targeting of human (beta)1- and (beta)2-adrenergic receptors following clathrin-mediated endocytosis. Journal of cell science. 2004;117(Pt 5):723–34.

33. Dehvari N, Hutchinson DS, Nevzorova J, Dallner OS, Sato M, Kocan M, et al. beta(2)-Adrenoceptors increase translocation of GLUT4 via GPCR kinase sites in the receptor C-terminal tail. Br J Pharmacol. 2012;165(5):1442–56.

34. Cannavo A, Koch WJ. Targeting beta3-Adrenergic Receptors in the Heart: Selective Agonism and beta-Blockade. J Cardiovasc Pharmacol. 2017;69(2):71–8.

35. Michel LYM, Farah C, Balligand JL. The Beta3 Adrenergic Receptor in Healthy and Pathological Cardiovascular Tissues. Cells-Basel. 2020;9(12).

36. Tuncay E, Olgar Y, Durak A, Degirmenci S, Bitirim CV, Turan B. beta(3) -adrenergic receptor activation plays an important role in the depressed myocardial contractility via both elevated levels of cellular free Zn(2+) and reactive nitrogen species. J Cell Physiol. 2019;234(8):13370–86.

37. Wright SC, Motso A, Koutsilieri S, Beusch CM, Sabatier P, Berghella A, et al. GLP-1R signaling neighborhoods associate with the susceptibility to adverse drug reactions of incretin mimetics. Nat Commun. 2023;14(1):6243.

38. Portha B, Chavey A, Movassat J. Early-life origins of type 2 diabetes: fetal programming of the beta-cell mass. Exp Diabetes Res. 2011;2011:105076.

39. Bitirim CV, Ozer ZB, Aydos D, Genc K, Demirsoy S, Akcali KC, et al. Cardioprotective effect of extracellular vesicles derived from ticagrelor-pretreated cardiomyocyte on hyperglycemic cardiomyocytes through alleviation of oxidative and endoplasmic reticulum stress. Sci Rep-Uk. 2022;12(1).

40. Lymperopoulos A. Clinical pharmacology of cardiac cyclic AMP in human heart failure: too much or too little? Expert Rev Clin Pharmacol. 2023;16(7):623–30.

41. Yildirim SS, Akman D, Catalucci D, Turan B. Relationship between downregulation of miRNAs and increase of oxidative stress in the development of diabetic cardiac dysfunction: junctin as a target protein of miR-1. Cell Biochem Biophys. 2013;67(3):1397–408.

42. Dincer UD, Bidasee KR, Guner S, Tay A, Ozcelikay AT, Altan VM. The effect of diabetes on expression of beta1-, beta2-, and beta3-adrenoreceptors in rat hearts. Diabetes. 2001;50(2):455-61.

43. Durak A, Turan B. Liraglutide provides cardioprotection through the recovery of mitochondrial dysfunction and oxidative stress in aging hearts. J Physiol Biochem. 2023;79(2):297–311.

44. Cheng T, Hao M, Takeda T, Bryant SH, Wang Y. Large-Scale Prediction of Drug-Target Interaction: a Data-Centric Review. AAPS J. 2017;19(5):1264–75.

45. Palve V, Liao Y, Remsing Rix LL, Rix U. Turning liabilities into opportunities: Off-target based drug repurposing in cancer. Semin Cancer Biol. 2021;68:209–29.

46. Rudmann DG. On-target and off-target-based toxicologic effects. Toxicol Pathol. 2013;41(2):310–4.

47. Taktaz F, Scisciola L, Fontanella RA, Pesapane A, Ghosh P, Franzese M, et al. Evidence that tirzepatide protects against diabetes-related cardiac damages. Cardiovasc Diabetol. 2024;23(1):112.

48. Graaf C, Donnelly D, Wootten D, Lau J, Sexton PM, Miller LJ, et al. Glucagon-Like Peptide-1 and Its Class B G Protein-Coupled Receptors: A Long March to Therapeutic Successes. Pharmacol Rev. 2016;68(4):954–1013.

49. Sonoda N, Imamura T, Yoshizaki T, Babendure JL, Lu JC, Olefsky JM. Beta-Arrestin-1 mediates glucagon-like peptide-1 signaling to insulin secretion in cultured pancreatic beta cells. Proceedings of the National Academy of Sciences of the United States of America. 2008;105(18):6614–9.

50. Jones B, Buenaventura T, Kanda N, Chabosseau P, Owen BM, Scott R, et al. Targeting GLP-1 receptor trafficking to improve agonist efficacy. Nat Commun. 2018;9(1):1602.

51. Pandey S, Mangmool S, Parichatikanond W. Multifaceted Roles of GLP-1 and Its Analogs: A Review on Molecular Mechanisms with a Cardiotherapeutic Perspective. Pharmaceuticals (Basel). 2023;16(6).

52. Cho YK, La Lee Y, Jung CH. The Cardiovascular Effect of Tirzepatide: A Glucagon-Like Peptide-1 and Glucose-Dependent Insulinotropic Polypeptide Dual Agonist. J Lipid Atheroscler. 2023;12(3):213–22.

53. Rowlands J, Heng J, Newsholme P, Carlessi R. Pleiotropic Effects of GLP-1 and Analogs on Cell Signaling, Metabolism, and Function. Front Endocrinol (Lausanne). 2018;9:672.

54. Rajan S, Dickson LM, Mathew E, Orr CM, Ellenbroek JH, Philipson LH, et al. Chronic hyperglycemia downregulates GLP-1 receptor signaling in pancreatic beta-cells via protein kinase A. Mol Metab. 2015;4(4):265–76.

55. Chen L, Chen X, Ruan B, Yang H, Yu Y. Tirzepatide protects against doxorubicin-induced cardiotoxicity by inhibiting oxidative stress and inflammation via PI3K/Akt signaling. Peptides. 2024;178:171245.

56. Huang JH, Chen YC, Lee TI, Kao YH, Chazo TF, Chen SA, et al. Glucagon-like peptide-1 regulates calcium homeostasis and electrophysiological activities of HL-1 cardiomyocytes. Peptides. 2016;78:91–8.

57. Ciminera AK, Shuck SC, Termini J. Elevated glucose increases genomic instability by inhibiting nucleotide excision repair. Life Sci Alliance. 2021;4(10).

58. Tanisha, Venkategowda S, Majumdar M. Amelioration of hyperglycemia and hyperlipidemia in a high-fat diet-fed mice by supplementation of a developed optimized polyherbal formulation. 3 Biotech. 2022;12(10):251.

59. Durak A, Akkus E, Canpolat AG, Tuncay E, Corapcioglu D, Turan B. Glucagon-like peptide-1 receptor agonist treatment of high carbohydrate intake-induced metabolic syndrome provides pleiotropic effects on cardiac dysfunction through alleviations in electrical and intracellular Ca(2+) abnormalities and mitochondrial dysfunction. Clin Exp Pharmacol Physiol. 2022;49(1):46–59.

60. Durak A, Olgar Y, Tuncay E, Karaomerlioglu I, Kayki Mutlu G, Arioglu Inan E, et al. Onset of decreased heart work is correlated with increased heart rate and shortened QT interval in high-carbohydrate fed overweight rats. Can J Physiol Pharmacol. 2017;95(11):1335–42.

61. Xu G, Kaneto H, Laybutt DR, Duvivier-Kali VF, Trivedi N, Suzuma K, et al. Downregulation of GLP-1 and GIP receptor expression by hyperglycemia: possible contribution to impaired incretin effects in diabetes. Diabetes. 2007;56(6):1551–8.

62. El K, Douros JD, Willard FS, Novikoff A, Sargsyan A, Perez-Tilve D, et al. The incretin co-agonist tirzepatide requires GIPR for hormone secretion from human islets. Nat Metab. 2023;5(6):945–54.

63. Samms RJ, Coghlan MP, Sloop KW. How May GIP Enhance the Therapeutic Efficacy of GLP-1? Trends in endocrinology and metabolism: TEM. 2020;31(6):410–21.

64. Coskun T, Sloop KW, Loghin C, Alsina-Fernandez J, Urva S, Bokvist KB, et al. LY3298176, a novel dual GIP and GLP-1 receptor agonist for the treatment of type 2 diabetes mellitus: From discovery to clinical proof of concept. Mol Metab. 2018;18:3–14.

65. Jovanovic A, Xu B, Zhu C, Ren D, Wang H, Krause-Hauch M, et al. Characterizing Adrenergic Regulation of Glucose Transporter 4-Mediated Glucose Uptake and Metabolism in the Heart. JACC Basic Transl Sci. 2023;8(6):638–55.

66. de Lucia C, Eguchi A, Koch WJ. New Insights in Cardiac beta-Adrenergic Signaling During Heart Failure and Aging. Front Pharmacol. 2018;9:904.

67. Moniotte S, Kobzik L, Feron O, Trochu JN, Gauthier C, Balligand JL. Upregulation of beta(3)-adrenoceptors and altered contractile response to inotropic amines in human failing myocardium. Circulation. 2001;103(12):1649–55.

68. Gauthier C, Rozec B, Manoury B, Balligand JL. Beta-3 adrenoceptors as new therapeutic targets for cardiovascular pathologies. Curr Heart Fail Rep. 2011;8(3):184–92.

69. Rozec B, Gauthier C. beta3-adrenoceptors in the cardiovascular system: putative roles in human pathologies. Pharmacol Ther. 2006;111(3):652–73.

70. Turan B, Tuncay E. Regulation of cardiac beta3-adrenergic receptors in hyperglycemia. Indian J Biochem Biophys. 2014;51(6):483–92.

