## Supplementary Document for "Dual GLP-1 and GIP receptor agonist Tirzapetide plays an off-target role in the modulation of the β-adrenoceptors and glucose metabolism in hyperglycemic or senescent cardiac cells"

**MTT assay for cell viability**

H9c2 cells were seeded on 96 well plates as 10000 cells/well. After overnight incubation, the cells were treated with TZPD (GlpBio, GC3840) at concentrations ranging between 20 nM - 300 nM for 24 h to optimize TZPD concentration. To optimize the antagonist doses, the cells were incubated with either GLP-1R antagonist (MedChem, HY-101116) ranging from 150 nM to 1.3 μM or GIP-R antagonist (MedChem, HY-P10138) ranging between 25 nM - 0.24 μM for 24 h. Following the incubation procedure, cell viability was evaluated by MTT cell viability assay (Bioworld, 42000092). Briefly, 0.5 mg of MTT solution was added per well. Following 3 h incubation, the supernatants were removed and 100 µL of DMSO was added to each well. A plate reader was used to measure signals at a 564 nm wavelength.

**Senescence-Associated β-galactosidase (SA-β-Gal) assay**

Mimicking senescence in H9c2 cells was detected by the Senescence β-Galactosidase Cell Staining Kit (Cell Signaling, 9860) in the SC group of H9c2 cells. A total of 10x10^5^ cells were seeded on 6 well plates and incubated with either 50 mg/mL D-galactose (SC group of cells) or 33 mM glucose (HC group of cells) for 48 h. Cells were incubated with 40 nM TZPD for 24 h during the incubation. A 1.3 µM of GLP-1R antagonist and 240 nM of GIP-R antagonists were added to the cell culture in the presence of TZPD and then the cells were incubated for 24 h. β-Galactosidase staining was performed according to the manufacturer’s instructions. The cells were visualized under the bright field microscope (Zeiss, PrimoVert).

**γH2A.X Staining**

Phosphorylated H2A.X (termed γH2A.X) staining in H9c2 cells was determined in NC, HG, D-gal (SC group of cells) group of H9c2 cells using anti- H2A.X (Thermo, MA5-14957 ) antibody using confocal microscopy (Zeiss LSM 980). Cells were incubated with 40 nM TZPD for 24 h during the incubation to obtain HG (33 mM glucose) + TZPD and SC+TZPD groups. To verify the effect of TZPD, the NC group of cells was incubated with insulin for 3 h. β-Actin (Santa Cruz, sc-47778) was used to identify the cellular structure. After fixation and permeabilisation of H9c2 cells with 4% paraformaldehyde and then 0.3% Triton-X100, the cells were incubated with primary antibody to monitor their localization on the membrane and cytoplasm. After overnight incubation of the cells, they were further incubated with appropriate secondary antibodies in the presence 0.25% Triton X-100/PBS for 5 min/each and incubated with Texas red goat anti-rabbit IgG (H&L) (1:500) (Rockland, 611-1902) and/or fluorescein goat anti-mouse IgG (H&L) (1:500) (Rockland, 610-1202) for 1 h at room temperature. A mounting Medium with DAPI (Novus Biologicals, H-1200-NB) was used for mounting.

**β_3_-ARs overexpression in H9c2 cell line**

We used β_3_-ARs overexpressed H9c2 cells (β3OE), previously obtained and used in our already published article (36). Briefly, the cell line was stably transfected with complementary DNA (cDNA) assembled in a lentiviral pCMV6‐Entry vector tagged with the C‐terminal Myc‐DDK tags (Origene, NM-013108, Origene, Rockville, MD). Following 2 weeks of all required procedures, the cells were harvested, and the protein level of β3‐AR was performed by Western blot analysis.

**Animals and ventricular cardiomyocytes to determine the cAMP and cGMP levels**

We used adult male Wistar rats either metabolic syndrome-induced (MetS) or non-induced (6-mo-old), as described previously (60). We also used elderly male rats (24-month-old) compared to those of adults as described previously (6). All animals were tested for validation of various parameters by measuring body weight, fasting blood glucose level, insulin level, HOMA‐IR, and oral glucose tolerance test (OGTT). The study was conducted under the Guide for the Care and Use of Laboratory Animals and approved by Ankara University (reference numbers 2015-12-137 and 2016-18-165). Freshly isolated cardiomyocytes were obtained by an enzymatic method in the left ventricle of the rats, as previously described (73).

***In silico* Analysis**

An *in silico* analysis was used to elucidate protein-protein interactions in the light of experimental data. Tirzepatide is a new peptide-based drug containing two non-essential amino acids. In this study, the interactions of Tirzepatide with β_1_-, β_2_-, and β_3_-adrenergic receptor (AR) proteins were analyzed *in silico* by using the protein-protein docking server; hddock (Huazhong University of Science and Technology, China) (71). The accuracy of the obtained data was compared with the data of the Human GLP-1R-Gs Complex (PDB ID:7FIM), whose structure is experimentally available.

PDB ID: 7FIM structure and the β_1_-AR (AF-P18090-F1), β_2_-AR (AF-P10608-F1), and β_3_-AR (AF-P26255-F1) structures obtained from the Alpha Fold site (EMBL-EBI, UK) (72) were analyzed on the hddock docking server. The energy values for the interactions of Tirzepatide with these proteins are calculated for comparison among proteins and given in Suppl. Table 2.

**Supplementary Figure Legends**

**Suppl. Fig. 1**. **(A)** The effect of tirzepatide (TZPD) on cell viability was determined by using the MTT assay. The H9c2 myoblasts were treated with various concentrations of TZPD (between 20 nM - 0.3 μM) for 24 h incubation. **(B)** Validation of senescence cells (SC group of cells) with detection of senescence-associated β-galactosidase (SA-β-Gal) in H9c2 cells treated with vehicle (NC group of cells), SC group, SC+1.3 μM GLP-1R antagonist +240 nM GIP-R antagonist (SC+ANT group), SC+TZPD group, SC+1.3 μM GLP-1R antagonist +240 nM GIP-R antagonist +TZPD (SC+ANT+TZPD group). **(C)** DNA double-strand breaks were detected by γH2AX staining in H9c2 cells treated with vehicle (NC group), SC group, SC+TZPD group, HG, HG+TZPD and NC+ Insulin (INS group). All experimental values of untreated cells (NC group cells) were set as 100% cell viability. Data are presented as mean±SEM. **p*<0.05 *vs.* NC group (*n*=3 independent experiments).

**Suppl. Fig. 2**. **(A)** Cell viability dose responses after treatment with GLP-1R (between 150 nM - 1.3 μM) or GIP-R (between 25 nM - 0.24 μM) antagonists for 24 h incubation. **(B)** The relative cAMP levels following the 30 min antagonist treatment in H9c2 cells in response to either 650 nM GLP-1R antagonist, 120 nM GIP-R antagonist, or the combination of 1.3 μM GLP-1R antagonist and 240 nM GIP-R antagonist. All experimental values of untreated cells (NC group cells) were set as 100% cell viability. Data are presented as mean±SEM. **p*<0.05 *vs.* NC group (*n*=3 independent experiments).

**Suppl. Fig. 3**. **In silico analysis of TZPD interactions with membrane receptors.** Human GLP and TZPD (the pink one) interaction **(A)** and in silico its experimental demonstration **(B)**. A 3D-representation of TZPD (the pink one) interaction with Human GIPR **(C)**. A 3D representation of TZPD (the pink one) and its interactions with β_1_-AR **(D)**, β_2_-AR **(E),** and β_3_-AR **(F)**, respectively.

**Suppl. Fig. 4**. The glucose consumption level was validated in **the** NC group in comparison to the INS-applied group. All experimental values of untreated cells (NC group cells) were set to 100% cell viability. Data are presented as mean±SEM. **p*<0.05 *vs.* NC group (*n*=3 independent experiments).

**Suppl. Fig. 5**. The cAMP levels in ventricular cardiomyocytes isolated freshly either metabolic syndrome (MetS) rats **(A)** or **(B)** elderly (Aged) rats compared to the normal adult rats (CON). The cGMP level was determined in the cardiomyocytes of elderly rats **(C)**. Data are presented as mean±SEM. **p*<0.05 *vs.* NC group (*n*=3 independent experiments).
