## Supplementary Table 1 for "Dual GLP-1 and GIP receptor agonist Tirzapetide plays an off-target role in the modulation of the β-adrenoceptors and glucose metabolism in hyperglycemic or senescent cardiac cells"

| **Gene** | **Forward** | **Reverse** |
| --- | --- | --- |
| β1-AR | GCTCACCAACCTCTTCATCA | CCGTCACACATAGCACGTCT |
| β2-AR | GGAATGACAGCGACTTCTTG | GGAATGACAGCGACTTCTTG |
| β3-AR | GGAATGACAGCGACTTCTTG | CAGGCTCCTTGCTAGATCTC |
| PKG | CAGGCTCCTTGCTAGATCTC | AAATCGGAATGAGCCCCCTG |
| GLP-1R | GTGCGAAGAGTCCAAGCAA | ATGGCTGAAGCGATGACCAA |
| GIP-R | CTACACGGGAAACCAGACCC | TCTGAGCGTCTCACAACCAC |
| IRS1 | CTACACCCGAGACGAACACT | TAACCTGCCAGACCTCCTTG |
| GLUT4 | GCTGTGAGTGAGTGCTTTCC | ACTGTTGTGTTCCTTTGCCC |

**Supplementary Table 1.**  Primers designed for qRT-PCR
