## Supplementary Table 2 for "Dual GLP-1 and GIP receptor agonist Tirzapetide plays an off-target role in the modulation of the β-adrenoceptors and glucose metabolism in hyperglycemic or senescent cardiac cells"

**Supplementary Table 2**. Binding energy levels were calculated as kcal/mol for the interaction of TZPD with the receptors.

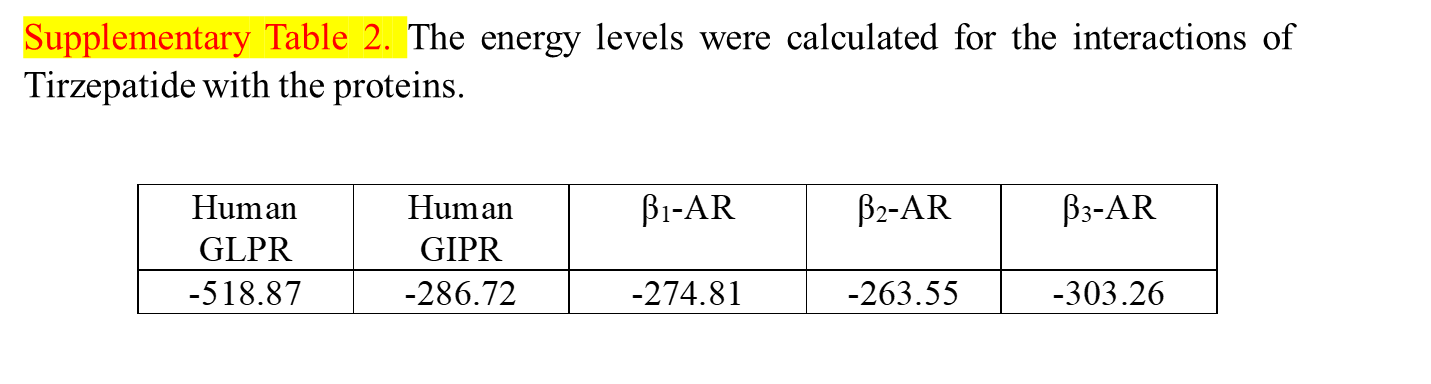
