## Supplementary figures and images for "Dual GLP-1 and GIP receptor agonist Tirzapetide plays an off-target role in the modulation of the β-adrenoceptors and glucose metabolism in hyperglycemic or senescent cardiac cells"

### Supplementary Figure 1

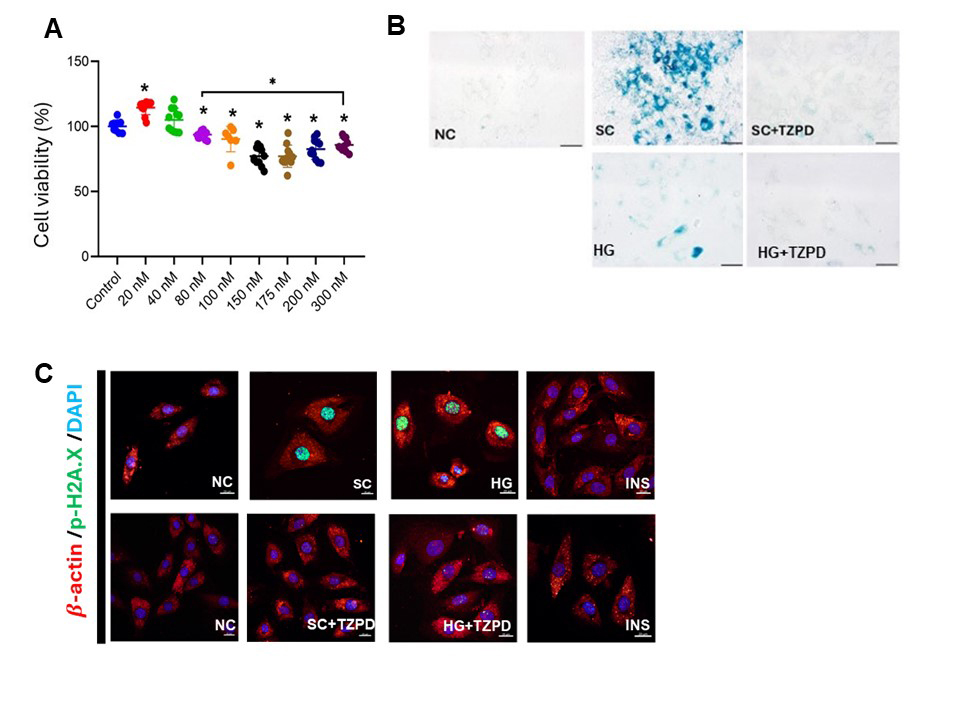

### Supplementary Figure 2

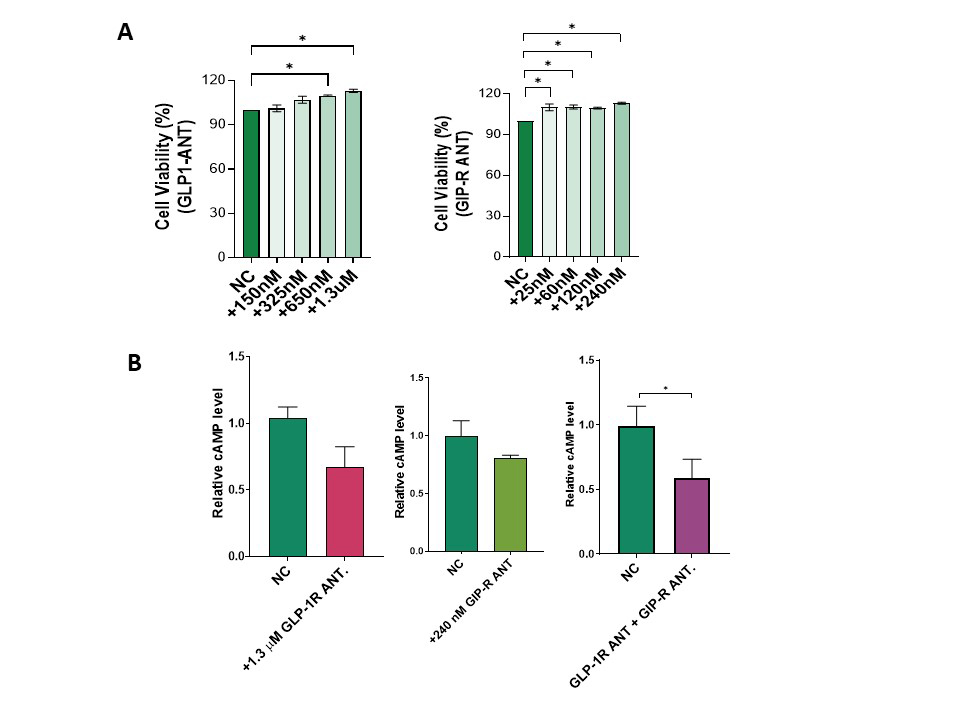

### Supplementary Figure 3

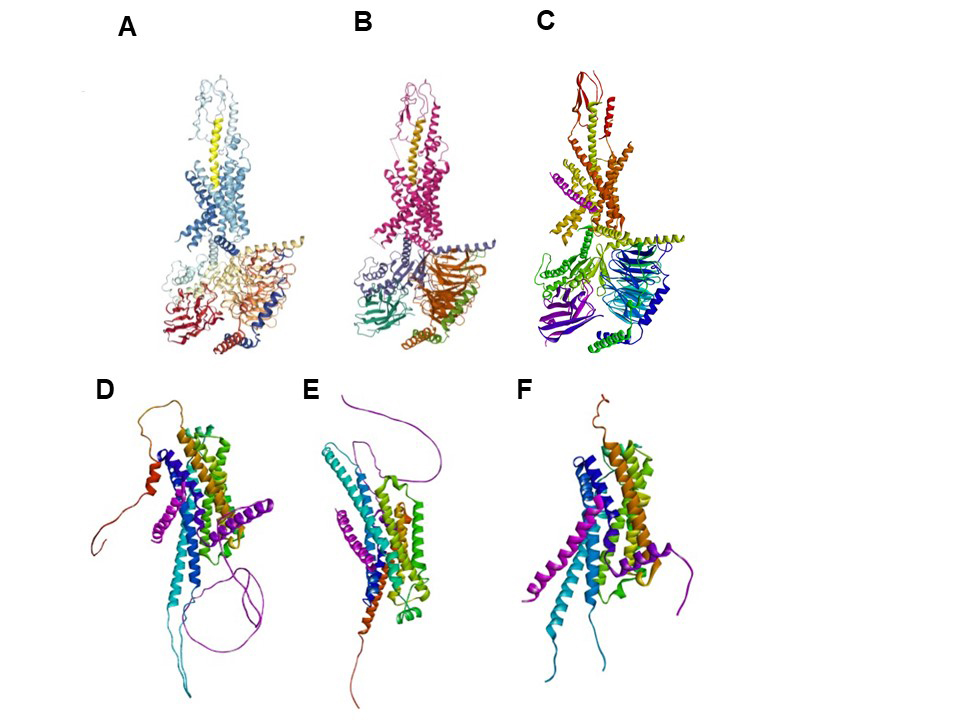

### Supplementary Figure 4

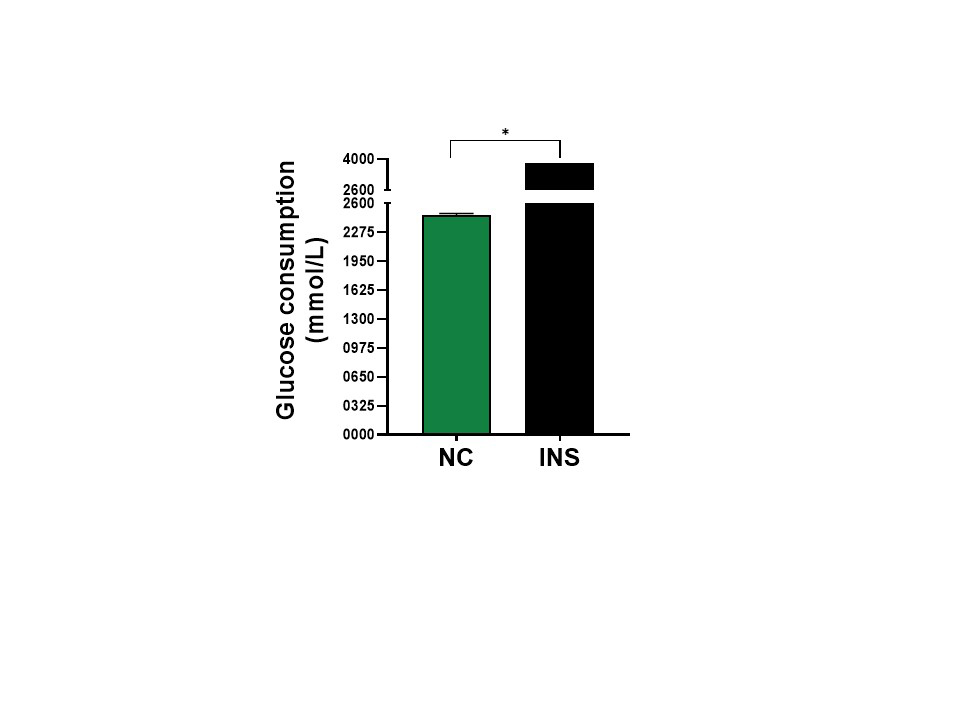

### Supplementary Figure 5

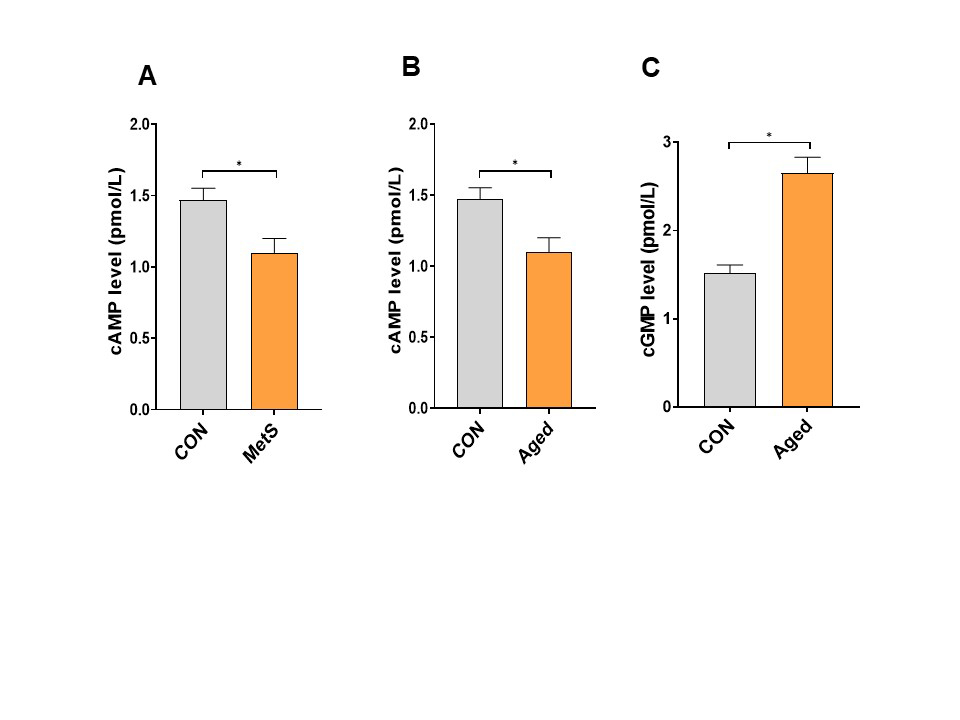
